# KCNQ2/3 Regulates the Synaptic Remodeling Associated with Drug Seeking in Addiction

**DOI:** 10.64898/2026.06.14.732187

**Authors:** Xinzhu Zhou, Xinjian Zhang, Yasuhiro Funahashi, Daisuke Tsuboi, Tetsuya Takano, Hisayoshi Kubota, Charles T. Yokoyama, Toshitaka Nabeshima, Kiyofumi Yamada, Kozo Kaibuchi, Taku Nagai

## Abstract

Drug-seeking behavior during withdrawal represents a critical obstacle to addiction treatment. In the nucleus accumbens, hyperactive dopamine D1 receptor-expressing medium spiny neurons (D1R-MSNs) promote cocaine-seeking through aberrant synaptic remodeling, including synapse formation and calcium-permeable AMPA receptor (CP-AMPAR) insertion. However, the intermediate molecular control mechanism remains unclear. We identified KCNQ2/3 potassium channels as key regulators of synaptic pathology during withdrawal. Cocaine-conditioned mice showed increased spine density, enhanced surface CP-AMPAR, and elevated neuronal activity 14 days after withdrawal. These phenotypes were reversed by repeated administration of KCNQ2/3 openers or D1R-MSN-specific expression of constitutively active KCNQ2. Functional restoration of KCNQ2/3 suppressed cocaine-seeking behavior and normalized D1R-MSN excitability. These findings suggest that sustained KCNQ2/3 deactivation drives synaptic remodeling during withdrawal, while channel activation offers a potential therapeutic strategy. Furthermore, the results position KCNQ2/3 as a master regulator of drug-seeking behavior and channel activation in D1R-MSNs as a logical target for relapse prevention in addiction.

## INTRODUCTION

Drug addiction is a chronic, relapsing brain disorder for which no effective pharmacological cure currently exists^1^. Despite extensive research, the medical community lacks universally recognized medications that can fundamentally prevent relapse and eliminate drug craving^2^. One of the greatest challenges in treating addiction is persistent drug-seeking behavior that emerges during the withdrawal period—a phase marked by heightened vulnerability to relapse even after prolonged abstinence^3^.

Addiction involves the nucleus accumbens (NAc)—a central brain region essential for reward processing, motivation, and memory^4^. The NAc comprises two main types of medium spiny neurons (MSNs): D1 dopamine receptor-expressing (D1R-MSNs), which are associated with reward signaling^5,6^, and D2 receptor-expressing (D2R-MSNs), which mediate aversive and inhibitory processes^6^. Chronic use of psychostimulants, such as cocaine, elevates extracellular dopamine in the NAc, increasing D1R-MSN activity and strengthening long-term maladaptive synaptic plasticity associated with craving and relapse^7,8^.

A hallmark of the withdrawal state is the formation of silent synapses—thin, immature spines lacking AMPA receptors^9^—and the subsequent insertion of high-conductance, calcium-permeable AMPA receptors (CP-AMPARs)^10^, particularly in D1R-MSNs^11^. This aberrant synaptic remodeling creates a permissive environment for drug seeking^12^. However, the upstream molecular mechanisms that regulate this spine remodeling remain elusive^13^.

In our previous work, we demonstrated that dopamine activates the PKA-Rap1-MAPK pathway in D1R-MSNs, leading to phosphorylation and functional inhibition of the voltage-gated potassium channel KCNQ2, thereby enhancing neuronal excitability and reward behaviors^14,15^. Based on these findings, we hypothesized that sustained deactivation of KCNQ2/3 channels in D1R-MSNs during the withdrawal period is a key driver of the maladaptive synaptic remodeling that underlies relapse.

In this study, we identified the KCNQ2/3 channel as a master regulator of synaptic and behavioral plasticity in cocaine withdrawal. Using pharmacological activation and viral genetic manipulation, we show that KCNQ2/3 activation reverses the formation of silent synapses, suppresses CP-AMPAR signaling, normalizes D1R-MSN excitability, and attenuates drug-seeking behavior. Together, our findings elucidate a fundamental mechanism of addiction-related synaptic pathology and suggest that KCNQ2/3 channels are promising target molecules for new therapeutic strategies on relapse prevention in cocaine addiction.

## RESULTS

### Repeated administration of a KCNQ2/3 potassium channel opener during the withdrawal period attenuates cocaine-seeking behavior

To investigate the effects of the KCNQ2/3 potassium channel opener on cocaine-seeking behavior triggered by contextual cues associated with the rewarding effects of cocaine, we conducted the conditioned place preference (CPP) test in mice. After cocaine conditioning, we administered intraperitoneal injections of the KCNQ2/3 potassium channel opener ICA-27243 to the mice once daily for seven consecutive days because a single injection of ICA-27243 had no effect (Figure S1). The control group received an equal volume of vehicle. To evaluate cocaine-seeking behavior during the cocaine withdrawal period, post-tests were carried out at 7 and 14 days after the last conditioning, namely cocaine withdrawal day 7 (WD7) and day 14 (WD14), respectively (Figure 1a). The CPP scores of the cocaine-conditioned group significantly increased compared with that of a saline-conditioned group on WD7, and also on WD14. Repeated treatment with ICA-27243 after cocaine conditioning significantly and dose-dependently decreased CPP scores in cocaine-conditioned mice on WD14, while ICA-27243 failed to decrease them on WD7 (Figure 1b). These results suggest that ICA-27243 ameliorates cocaine-seeking behavior, albeit with delayed onset.

**Figure 1.**
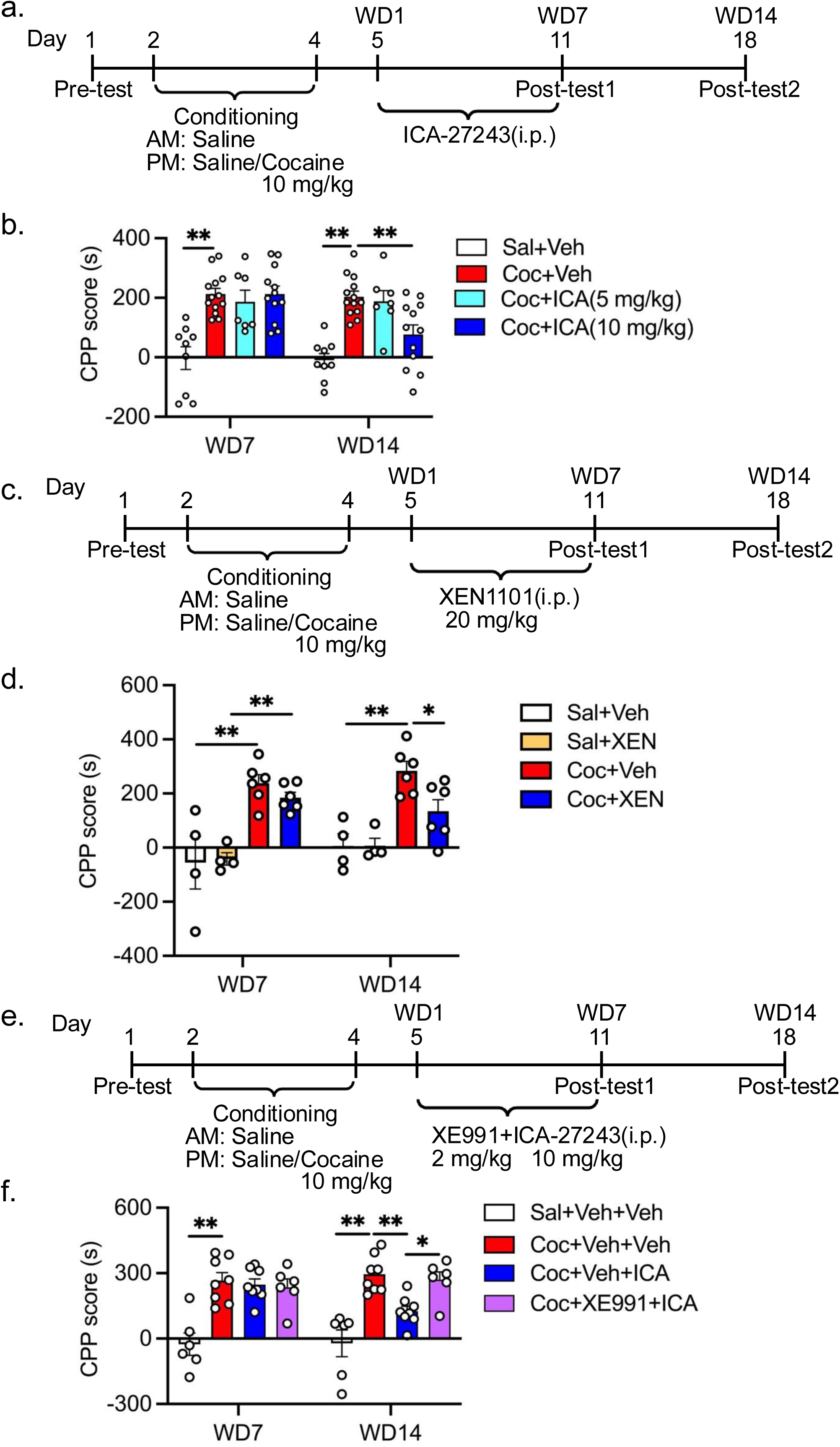
KCNQ2/3 potassium channel opener attenuates cocaine-seeking behavior after 14-day withdrawal a,b. Effect of ICA-27243 on cocaine-induced CPP after 14-day withdrawal. a. Timeline of CPP procedure. Mice were administered ICA-27243 or vehicle intraperitoneally for seven days, commencing on the first day of cocaine withdrawal (WD1). Post-tests were conducted on WD7 (day 11) and WD14 (day 18). b. The CPP score of WD7 and WD14. The administration of ICA-27243 (10 mg/kg) resulted in the attenuation of the CPP score induced by cocaine on WD14. Two-way ANOVA: Drug effect, F (3, 37)=14.48, P<0.01; time effect, F(1, 37)=6.899, P<0.05; interaction effect, F (3, 37)=6.026, P<0.01; Tukey’s post-hoc test: **p<0.01. Saline+Vehicle group (n=9), Cocaine+Vehicle group (n=13), Cocaine+ICA-27243 (5mg/kg) group (n=7), Cocaine+ICA-27243(10mg/kg) group (n=12). **c,d.** Effect of XEN1101 on cocaine-seeking behavior after 14-day withdrawal. c. Timeline of CPP procedure. Mice were administered XEN1101 or vehicle intraperitoneally for 7 days, commencing on the first day of cocaine withdrawal (WD1). Post-tests were conducted on WD7 (day 11) and WD14 (day 18). d. The CPP score of WD7 and WD14. The administration of XEN1101 (20 mg/kg) resulted in the attenuation of the CPP score induced by cocaine on WD14. Two-way ANOVA: Drug effect, F(3, 16)=20.31, P<0.01; time effect, F(1, 16)=0.7906, P=0.3871; interaction effect, F(3, 16)=0.8572, P=0.4832; Tukey’s post-hoc test: **p<0.01. Saline+Vehicle group (n=4), Saline+XEN1101 (20mg/kg) group (n=4), Cocaine+Vehicle group (n=6), Cocaine+XEN1101 (20mg/kg) group (n=6). **e, f.** Effect of XE991 on ameliorating the effect of ICA-27243 on cocaine-induced CPP after 14-day withdrawal. e. Timeline of CPP procedure. Mice were administered ICA-27243(10mg/kg), vehicle and/or XE991 (2mg/kg) intraperitoneally for seven days, commencing on the first day of cocaine withdrawal (WD1). Post-tests were conducted on WD7 (day 11) and WD14 (day 18). Mice were administered XE991 5 min before the ICA-27243 treatment. f. CPP score of WD7 and WD14. The concomitant administration of XE991 suppressed the attenuating effect of ICA-27243 on cocaine-seeking behavior after 14-day withdrawal. Two-way ANOVA: Drug effect, F(3, 24)=14.71, P<0.01; time effect, F(1, 24)=0.8858, P=0.3560; interaction effect, F(3, 24)=7.243, P<0.01; Tukey’s post-hoc test; *p<0.05, **p<0.01. Saline+Vehicle+Vehicle group (n=6), Cocaine+Vehicle+Vehicle group (n=8), Cocaine+Vehicle+ICA-27243 group (n=8), Cocaine+XE991+ICA-27243 group (n=6).

Next, we examined the effect of other KCNQ2/3 potassium channel openers, such as XEN1101 and ICA-110381, on cocaine-induced CPP in mice (Figures 1c,d and S2a,b). Consistent with the result of ICA-27243, repeated treatment of both XEN1101 or ICA-110381 significantly decreased CPP scores in cocaine-conditioned mice on WD14 (Figures 1d and S2b).

To confirm that the ameliorating effect of ICA-27243 on cocaine-seeking behavior is associated with the activation of KCNQ2/3 channels, we used the KCNQ2/3 potassium channel blocker XE991. XE991 (2mg/kg, i.p.) was administered to mice 5 min before the injection of ICA-27243 (Figure 1e). Co-administration of XE991 inhibited the ameliorating effect of ICA-27243 on cocaine-induced CPP (Figure 1f).

These results suggest that pharmacological activation of the KCNQ2/3 potassium channel attenuates cocaine-seeking behavior during withdrawal.

### Activation of KCNQ2 in D1R-MSNs of the NAc attenuates cocaine-seeking behavior

The specific brain regions and cell types mediating the ameliorating effect of KCNQ2/3 channel openers on cocaine-seeking behavior remain unclear. Previous evidence confirms that the NAc plays a key role in cocaine addiction, and D1R-MSNs play a more important role compared with D2R-MSNs^16,17^. To examine if the KCNQ2/3 potassium channel opener attenuates cocaine-seeking behavior mediated by accumbal D1R-MSNs, we manipulated KCNQ2 channel activity specifically in D1R-MSNs by over-expressing a constitutively active form of KCNQ2 (KCNQ2-CA)^18^. AAV-CW3SL-Flex-KCNQ2-CA was microinjected into the NAc of Drd1-Cre mice (Figure 2a,b). The function of KCNQ2-CA was validated by its expression in D1R-MSNs, to assess whether it is able to inhibit kainic acid-induced seizures and c-Fos expression in D1R-MSNs (Figure S3). KCNQ2-CA expression in the NAc was also confirmed after CPP conditioning (Figure 2c). Cocaine-conditioned mice expressing mCherry in accumbal D1R-MSNs showed increased CPP scores compared with saline-conditioned mice (Figure 2d). Consistent with the effects of repeated administration of the KCNQ2/3 potassium channel opener, cocaine-conditioned mice over-expressing KCNQ2-CA in accumbal D1R-MSNs showed a significant reduction in CPP score compared with cocaine-conditioned mice expressing mCherry, as observed on WD14 (Figure 2d).

**Figure 2.**
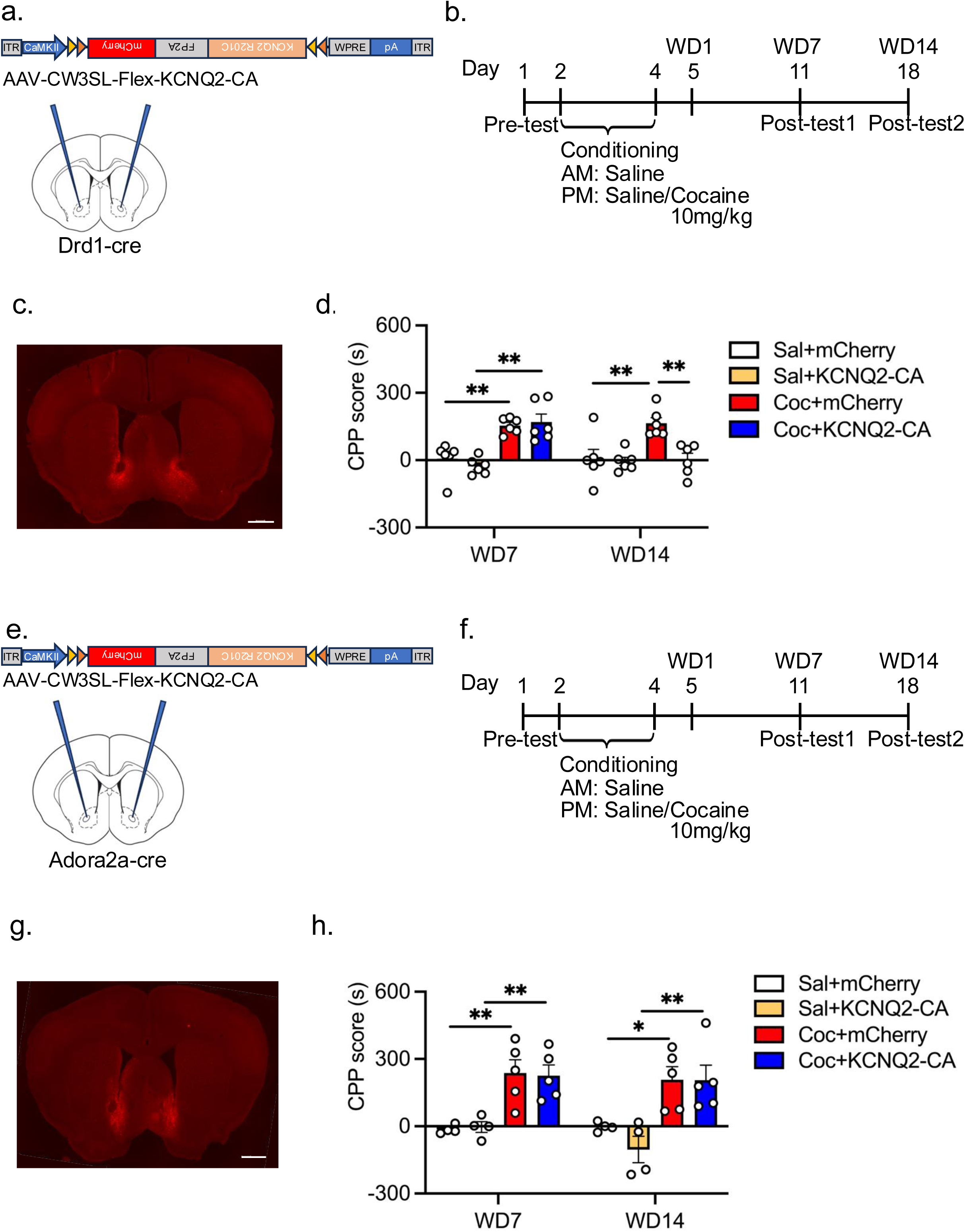
KCNQ2-CA over-expression in D1R-MSNs attenuates cocaine-seeking behavior after 14-day withdrawal a−d. Effect of KCNQ2-CA in D1-MSNs on cocaine-induced CPP after 14-day withdrawal. a. Schematic of AAV-mediated KCNQ2-CA expression in accumbal D1R-MSNs. Using stereotaxic injections, AAV-KCNQ2-CA (5 x 10^12^ copies/ml) was bilaterally injected into the NAc of mice. The diagram shows the AAV constructs and stereotaxic injection of AAVs into the NAc of Drd1-Cre transgenic mice. b. Timeline of experimental procedure. Sixteen days after the AAV injection, the mice were subjected to a CPP experiment. c. Representative coronal brain section showing the expression of mCherry five weeks after AAV injection into the NAc. d. CPP score of WD7 and WD14. Two-way ANOVA: Virus effect, F(3,20)=11.11, P<0.01; time effect, F(1,20)=5.695, P<0.05; interaction effect, F(3,20)=9.713, P<0.01; Tukey’s post-hoc test; *p<0.05, **p<0.01. n=6. **e−h.** Effect of KCNQ2-CA in D2R-MSNs on cocaine-induced CPP after 14-day withdrawal. e. Schematic of AAV-mediated KCNQ2-CA expression in accumbal D2R-MSNs. Using stereotaxic injections, AAV-KCNQ2-CA (5 x 10^12^ copies/ml) was bilaterally injected into the NAc of mice. The diagram shows the AAV constructs and stereotaxic injection of AAVs into the NAc of Adora2a-Cre transgenic mice. f. Timeline of experimental procedure. Sixteen days after the AAV injection, the mice were subjected to a CPP experiment. g. Representative coronal brain section showing the expression of mCherry five weeks after AAV injection into the NAc. h. CPP score of WD7 and WD14. Two-way ANOVA: Virus effect, F(3,14)=14.12, P<0.01; time effect, F(1,14)=1.056, P=0.3216; interaction effect, F(3,14)=0.4554, P=0.7177; Tukey’s post-hoc test; *p<0.05, **p<0.01. Saline-conditioned mice (n=4), cocaine-conditioned mice (n=5).

To manipulate KCNQ2 channels in D2R-MSNs, we microinjected AAV-CW3SL-Flex-KCNQ2-CA into the NAc of Adora2a-Cre mice (Figure 2e,g). The CPP experiment commenced 16 days after the microinjection (Figure 2f). The CPP scores of cocaine-conditioned mice over-expressing KCNQ2-CA in accumbal D2R-MSNs were comparable to those of cocaine-conditioned mice expressing mCherry on WD14 (Figure 2h).

These results strongly indicate that KCNQ2 in the accumbal D1R-MSNs, but not D2R-MSNs, is responsible for the amelioration of cocaine-seeking behavior.

### KCNQ2/3 potassium channel opener attenuates the cocaine-induced increase in spine density of D1R-MSNs

Dendritic spines are critical structural determinants of addiction vulnerability^19^. A previous study showed that cocaine administration induces morphological changes in dendritic spines of the NAc^20^. To investigate if the KCNQ2/3 potassium channel opener attenuates the changes in dendritic spine morphology as well as cocaine-seeking behavior, we microinjected AAV-Sp-Cre and AAV-Flex-EGFP into the NAc of C57BL/6 mice to label spines of accumbal D1R-MSNs (Figure 3a). We carried out the CPP experiment two weeks after virus injection (Figure 3b). Mice were sacrificed on WD1 (without ICA-27243 injection), WD7 (immediately after post-test 1), or WD14 (immediately after post-test 2) to analyze spine morphology in D1R-MSNs of the NAc (Figure 3b). Compared with the saline-conditioned control group, cocaine-conditioned mice showed a significant increase in total spine density in D1R-MSNs of the NAc on WD1, WD7, and WD14 (Figure 3c,d). Interestingly, there was a marked increase in total spine density on WD14, indicating a progressive elevation of spine density across the cocaine withdrawal period. We examined the effect of ICA-27243 administration on spine density of D1R-MSNs and found that spine density in ICA-27243-treated cocaine-conditioned mice was comparable to that of vehicle-treated cocaine-conditioned mice on WD7 (Figure 3c,d). However, the spine density in ICA-27243 treated cocaine-conditioned mice was significantly reduced to the level of vehicle-treated cocaine-conditioned mice on WD14 (Figure 3c,d). ICA-27243 itself had no effect on spine density in saline-treated mice on WD7 and WD14 (Figure 3c,d). We also analyzed spine subtypes and found that the overall increase in spine density of cocaine-conditioned mice was primarily due to an increase in thin spines (Figure S4a), with a slight increase in stubby and mushroom spines (Figure S4b,c). Repeated ICA-27243 administration reduced the number of thin, stubby, and mushroom spines in cocaine-conditioned mice on WD14 (Figure S4). Repeated ICA-27243 administration itself had no effect on spine density of all subtypes in saline-treated mice on WD7 and WD14 (Figure S4).

**Figure 3.**
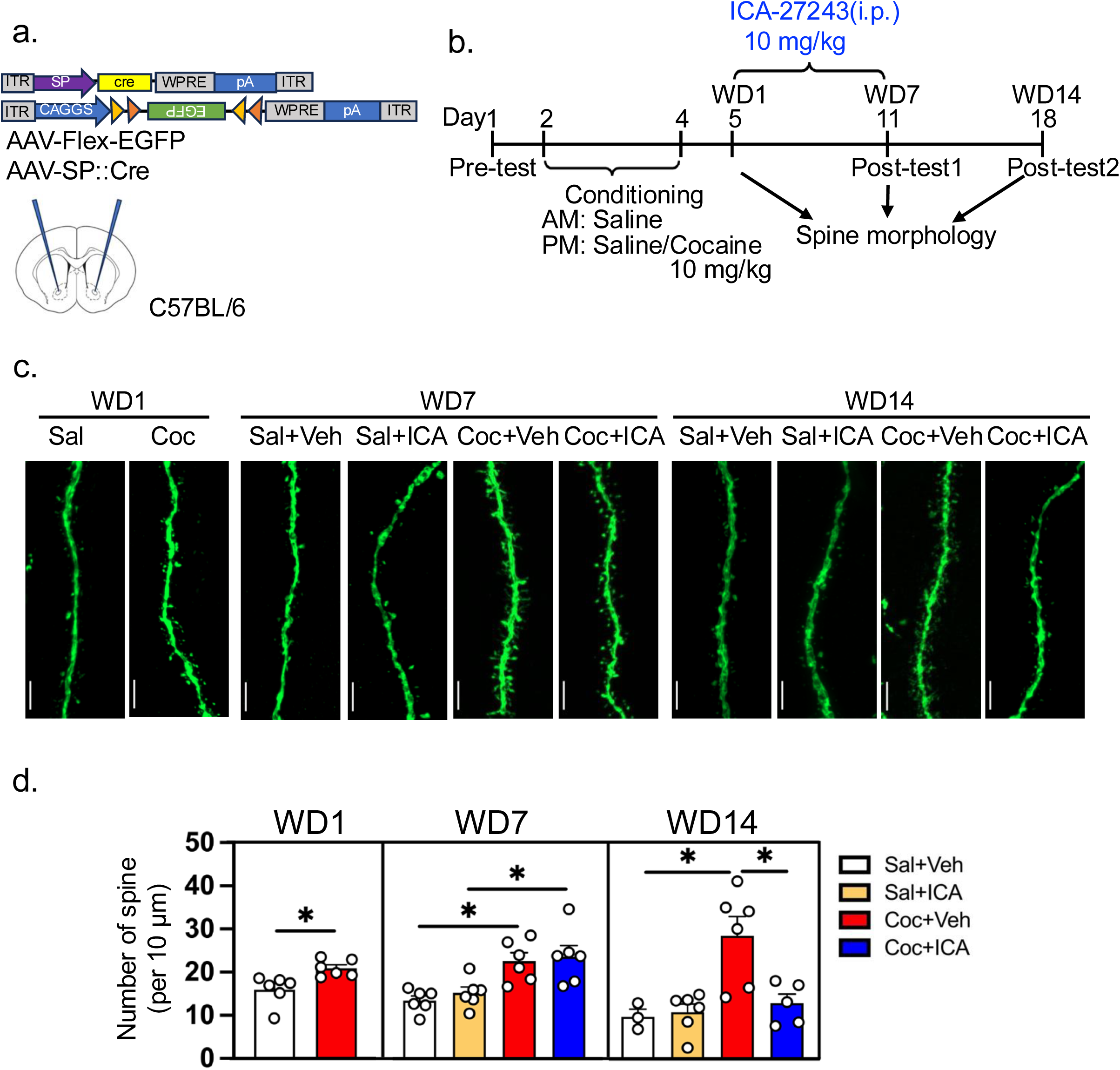
KCNQ2/3 potassium channel opener attenuates the cocaine-induced increase in spine density in D1R-MSNs. **a.** Schematic of AAV-mediated EGFP expression in accumbal D1R-MSNs. Using stereotaxic injections, AAV-Flex-EGFP and AAV-SP-Cre were bilaterally injected into the NAc of mice. The diagram shows the AAV constructs and stereotaxic injection of AAVs into the NAc of Drd1-Cre transgenic mice. **b.** Timeline of CPP procedure. Mice were administered ICA-27243 or vehicle intraperitoneally for seven days, commencing on the first day of cocaine withdrawal (WD1). Post-tests were conducted on WD7 (day 11) and WD14 (day 18). Spine analysis was carried out on WD1, WD7, and WD14. **c.** Representative image of viral-mediated EGFP expression in the secondary dendrites of D1R-MSNs. The scale bar indicates 5μm. **d.** Quantification of spine density on WD1, WD7, and WD14. Two-tailed unpaired t-test; WD1, t=3.062, P<0.05. Two-way ANOVA; WD7, ICA-27243, F(1,20)=0.5484, P=0.4676; cocaine, F(1,20)=22.67, P<0.01; interaction: F(1,20)=0.04399, P=0.8360; WD14, ICA-27243, F(1,20)=9.175, P<0.01; cocaine, F(1,20)=14.9, P<0.01; interaction, F(1,20)=12.57, P<0.01. Tukey’s post-hoc test; *p<0.05.

These results are consistent with the result of the CPP test in the present study (Figure 1b), indicating that the KCNQ2 opener may attenuate cocaine-seeking behavior by inhibiting cocaine-induced structural plasticity of dendritic spines in the accumbal D1R-MSNs.

### KCNQ2/3 potassium channel opener attenuates cocaine-seeking behavior by suppressing the cell surface expression of CP-AMPARs in D1R-MSNs

CP-AMPARs are a subtype of AMPA-type glutamate receptors that lack the GluA2 subunit^10,21,22^. Previous studies have reported increased CP-AMPAR expression after cocaine withdrawal^23,24^. The increase in CP-AMPARs is closely related to synaptic plasticity^10,25^, which can lead to enhanced sensitivity of neurons to excitatory input, thereby strengthening drug-related memories and behaviors^12^. The significant increase of spine density in accumbal D1R-MSNs was observed after 14 days of withdrawal following cocaine conditioning. Next, the expression of CP-AMPAR was investigated in D1R-MSNs of the NAc after withdrawal, with a particular focus on their surface membrane expression, using TurboID biotin labeling technology. AAV-Ef1a-DIO-TurboID-surface was injected into the NAc of Drd1-Cre mice to label membrane protein of D1R-MSNs (Figure 4a), and a CPP experiment was performed three weeks later. After post-test 2, mice were administered biotin, and the biotinylated surface membrane proteins in the NAc were collected by avidin agarose beads (Figure 4b). Immunoblot analysis revealed that the membrane surface expression of AMPA GluA1 and GluA3 subunits in the accumbal D1R-MSNs significantly increased in cocaine-conditioned mice on WD14 (Figure 4c,d,g,h). Repeated ICA-27243 administration reduced the increased membrane surface expression of AMPA GluA1 and GluA3 subunits in the NAc of cocaine-conditioned mice (Figure 4c,d,g,h). The membrane surface expression of GluA2 was comparable in the cocaine-conditioned or saline-conditioned groups with or without repeated ICA-27243 administration (Figure 4e,f). No changes were found in the total expression levels of GluA1, GluA2, and GluA3 among all mice (Figure 4c−h).

**Figure 4.**
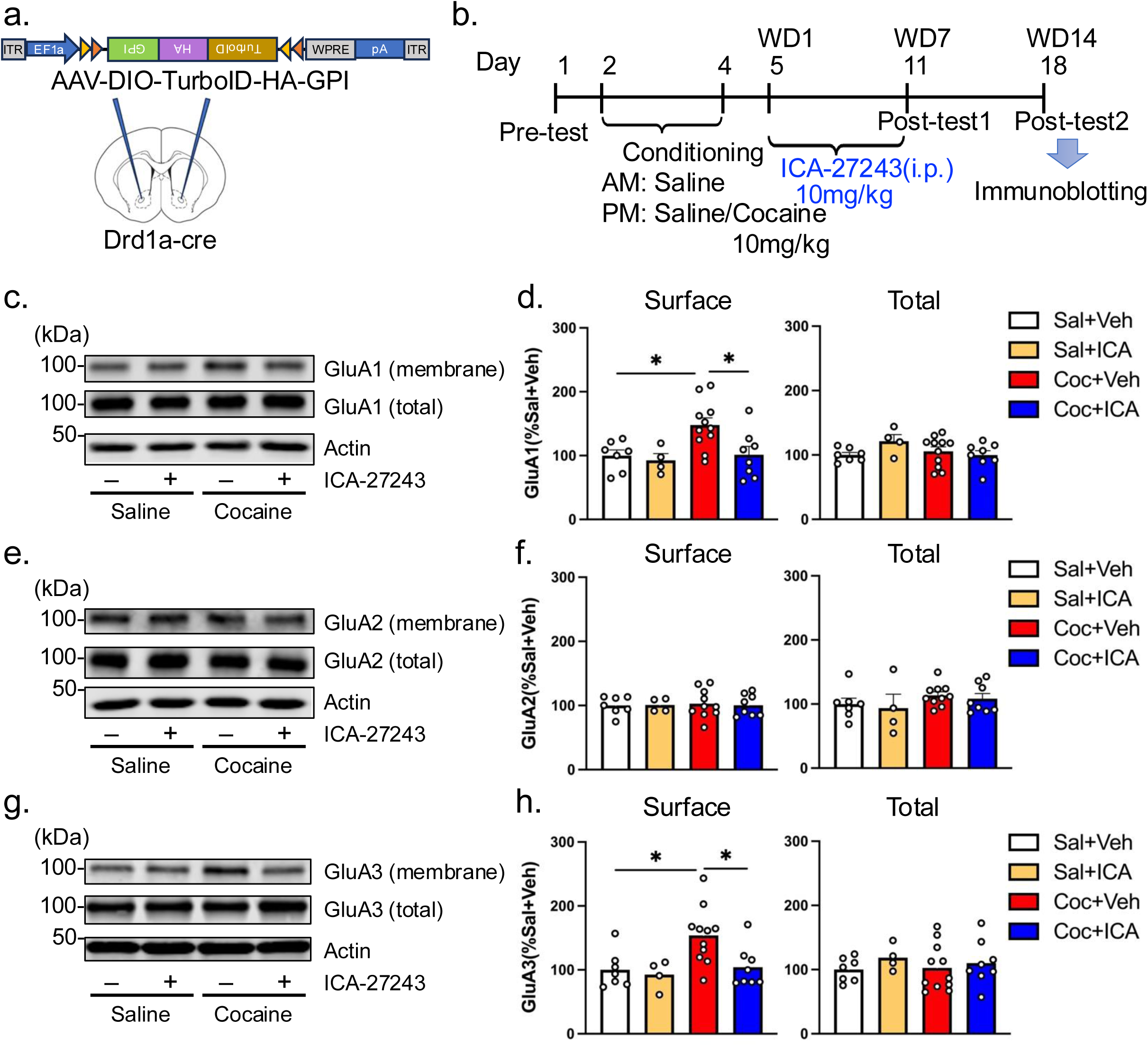
KCNQ2/3 potassium channel opener attenuates cocaine-seeking behavior by suppressing the cell surface expression of calcium-permeable AMPA receptors in D1R-MSNs. **a.** Schematic of AAV-mediated TurboID-surface expression in accumbal D1R-MSNs. Using stereotaxic injections, AAV-Flex-TurboID-surface (1 x 10^12^ copies/ml) was bilaterally injected into the NAc of mice. The diagram shows the AAV constructs and stereotaxic injection of AAVs into the NAc of Drd1-Cre transgenic mice. **b.** Timeline of experimental procedure. Three weeks after AAV-Flex-TurboID-surface injection, mice were administered ICA-27243 (10mg/kg) or vehicle intraperitoneally for seven days, commencing on the first day of cocaine withdrawal (WD1). Post-tests were conducted on WD7 (day 11) and WD14 (day 18). After the CPP test on WD14, biotin was immediately injected into the mice, and 1 h later, the mice were euthanized and the NAc was extracted. Expression level of calcium-permeable AMPA receptor was analyzed by immunoblotting. **c.** Representative image of GluA1 immunoblot analysis. **d.** Quantification for expression level of GluA1 subunit on cell surface (left panel). Two-way ANOVA: Cocaine, F(1, 26)=4.384, P<0.05; ICA-27243, F(1, 26)=4.851, P<0.05; interaction, F(1, 26)=2.287, P=0.1425. Quantification for expression level of GluA1 subunit in D1R-MSNs (right panel). Two-way ANOVA: Cocaine, F(1, 26)=1.085, P=0.3072; ICA-27243, F(1, 26)=1.097, P=0.3046; interaction, F(1, 26)=3.167, P=0.0868. Tukey’s post-hoc test; *p<0.05. **e.** Representative image of GluA2 immunoblot analysis. **f.** Quantification for expression level of GluA2 subunit on cell surface (left panel). Two-way ANOVA: Cocaine, F(1, 25)=0.01375, P=0.9076; ICA-27243, F(1, 25)=0.03266, P=0.8580; interaction, F(1, 25)=0.05675, P=0.8136. Quantification for expression level of GluA2 subunit in D1R-MSNs (right panel). Two-way ANOVA: Cocaine, F(1, 25)=0.3558, P=0.5562; ICA-27243, F(1, 25)=2.114, P=0.1584; interaction, F(1, 25)=0.002871, P=0.9577. Tukey’s post- hoc test. **g.** Representative image of GluA3 immunoblot analysis. **h.** Quantification for expression level of GluA3 subunit on cell surface (left panel). Two-way ANOVA: Cocaine, F(1, 26)=5.510, P<0.05; ICA-27243, F(1, 26)=4.240, P<0.05; interaction, F(1, 26)=2.323, P=0.1396. Quantification for expression level of GluA3 subunit in D1R-MSNs (right panel). Two-way ANOVA: Cocaine, F(1, 26)=0.05526, P=0.8160; ICA-27243, F(1, 26)=1.165, P=0.2904; interaction, F(1, 26)=0.2127, P=0.6485. Tukey’s post-hoc test; *p<0.05.

These results suggest that the membrane surface content, but not total protein amount, of CP-AMPARs in the accumbal D1R-MSNs increased during cocaine withdrawal, and repeated administration of the KCNQ2/3 channel opener can suppress membrane surface CP-AMPAR content.

### KCNQ2/3 potassium channel opener attenuates the cocaine-induced enhancement of AMPA receptor response in the NAc

Stimulation of CP-AMPAR can increase permeability of calcium ions followed by activation of CaMKII signaling^26^. To examine the effect of a KCNQ2/3 potassium channel opener on CP-AMPAR signaling in the NAc of cocaine-conditioned mice, we prepared acute striatal slices (including the NAc) of the mice after post-test 2 at WD14. The striatal slices were treated with an AMPAR agonist, (S)-AMPA (10 μM). The phosphorylation levels of AMPA GluA1 subunit (S831) and NMDA receptor NR2B subunit (S1303) that are substrates of CP-AMPAR/Ca^2+^/CaMKII signaling were analyzed by immunoblotting (Figure 5a). In saline-conditioned mice, the phosphorylation level of GluA1 slightly, but not significantly, increased in response to (S)-AMPA, indicating that CP-AMPAR expression is minimal (Figure 5b,c). (S)-AMPA treatment markedly and significantly promoted the phosphorylation level of GluA1 in cocaine-conditioned mice compared with that in the vehicle-treated control group, suggesting a hyperactivity of AMPAR signaling in cocaine-conditioned mice on WD14 (Figure 5b,c). The hyperactivation of AMPAR signaling in cocaine-conditioned mice was significantly reduced with repeated ICA-27243 administration. ICA-27243 treatment had no effect on the phosphorylation level of GluA1 in saline-conditioned mice (Figure 5b,c). Similar results were observed in the phosphorylation levels of NMDA receptor NR2B subunit (Figure 5d,e).

**Figure 5.**
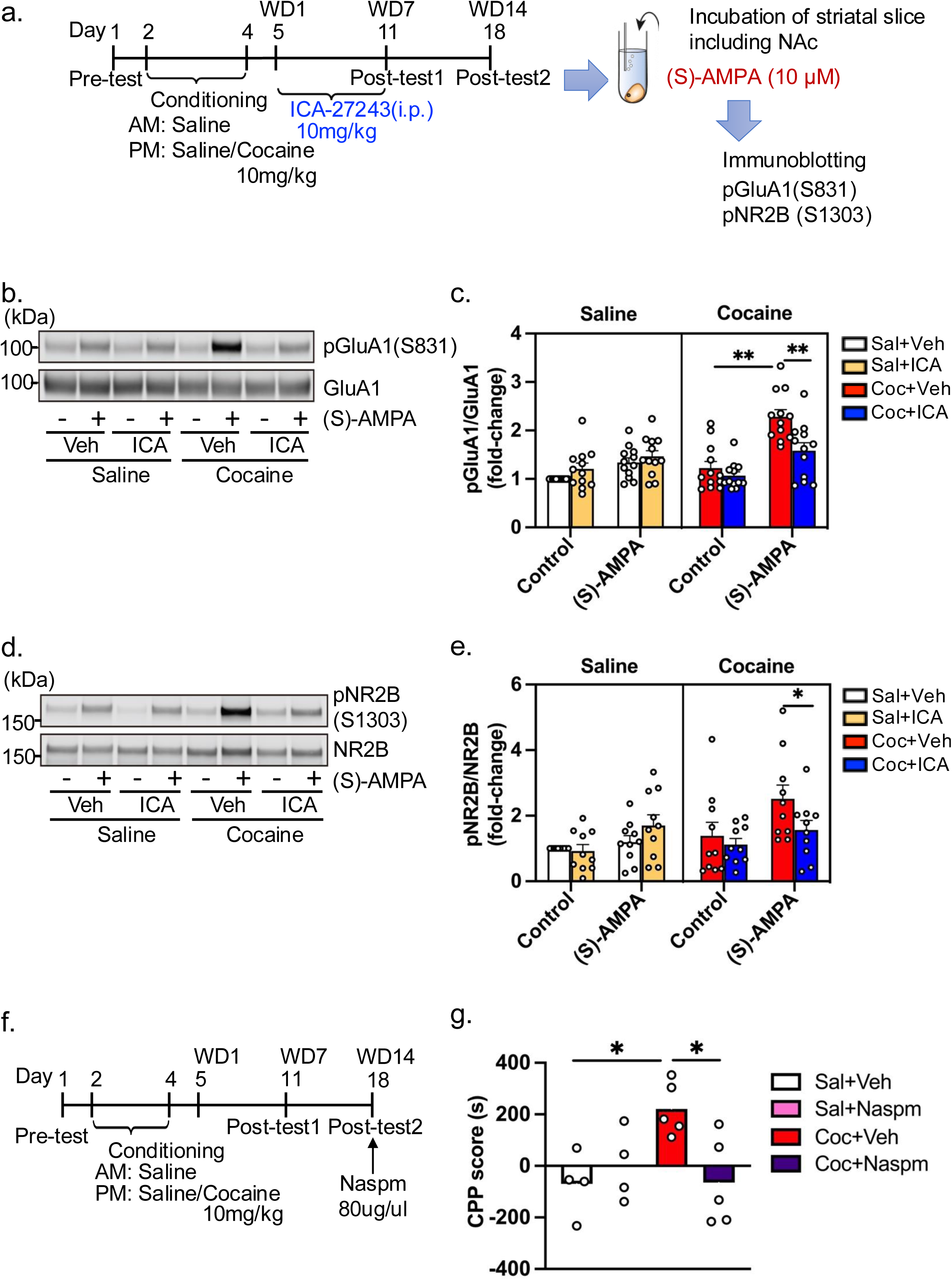
KCNQ2/3 potassium channel opener attenuates the cocaine-induced enhancement of AMPA receptor responses in the NAc. **a.** Timeline of experimental procedure. Mice were administered ICA-27243 or vehicle intraperitoneally for seven days from the first day of cocaine withdrawal (WD1). Post-tests were conducted on WD7 (day 11) and WD14 (day 18). Striatal slices including the NAc of mice were prepared after the CPP test on WD14. The slices were treated with (S)-AMPA (10 μM) for 15 s. Phosphorylation levels of AMPA receptor GluA1 subunit (S831) and NMDA receptor NR2B subunit (S1303) were analyzed by immunoblotting. **b,c.** Phosphorylation level of AMPA receptor GluA1 subunit (S831). **b.** Representative image of immunoblot analysis. **c.** Quantification of phosphorylation level of GluA1 subunit (S831). Three-way ANOVA: Cocaine, F(1, 44)=11.06, P<0.01; ICA-27243, F(1, 44)=2.702, P=0.1074; interaction, F(1, 44)=13.98, P=0.01. Tukey’s post-hoc test; **p<0.01. **d,e.** Phosphorylation level of NMDA receptor NR2B subunit (S1303). **d.** Representative image of immunoblot analysis. **e.** Quantification of phosphorylation level of NR2B subunit (S1303). Three-way ANOVA: Cocaine, F(1, 36)=3.229, P=0.0807; ICA-27243, F(1, 36)=1.897, P=0.1769; interaction, F(1, 36)=8.313, P=0.0066. Tukey’s post-hoc test; *p<0.05. **f,g.** Effects of Naspm (CP-AMPA-R antagonist; 80ug/ul) on cocaine-seeking behavior after 14-day withdrawal. **f.** Timeline of experiment procedure. Mice were microinjected with Naspm or vehicle into the NAc 10 min before the CPP test on WD14. **g.** CPP score of WD7 and WD14. Three-way ANOVA: Drug, F(3, 14)=7.714, P<0.01; WD, F(1, 14)=1.878, P=0.1922; interaction, F(3, 14)=3.695, P<0.05. Tukey’s post-hoc test, *p<0.05. Saline+Vehicle group (n=4), Cocaine+Vehicle group (n=5), Saline+Naspm group (n=4), Cocaine+Naspm group (n=5).

These results indicate that 14-day withdrawal after cocaine conditioning enhances CP-AMPAR signaling in mice, while repeated administration of ICA-27243 can block hyperphosphorylation through CP-AMPARs in cocaine-conditioned mice.

To confirm the role of accumbal CP-AMPARs in cocaine-seeking behavior, we microinjected a CP-AMPAR antagonist Naspm (80 μg/μl/site) into the NAc of saline or cocaine-conditioned mice 15 min before post-test 2 on WD14 (Figure 5f). Microinjection of Naspm into the NAc significantly reduced the CPP scores of cocaine-conditioned mice compared with those of the vehicle group. However, the CPP scores were comparable between vehicle- and Naspm-microinjected groups in saline-conditioned mice (Figure 5g).

These results suggest that the activation of CP-AMPAR signaling during cocaine withdrawal leads to cocaine-seeking behavior, while KCNQ2/3 channel opener administration attenuates the behavior through the suppression of CP-AMPAR signaling.

### KCNQ2/3 potassium channel opener attenuates the abnormal activity of accumbal D1R-MSNs in cocaine-seeking behavior

Since D1R-MSNs play a crucial role in cocaine-seeking behavior, we investigated the neuronal activity of D1R-MSNs in the NAc of cocaine-conditioned mice after repeated administration of ICA-27243.

We carried out the CPP experiment using Drd1-mVenus transgenic mice, in which D1R-MSNs express a variant of yellow fluorescent protein (mVenus). We collected cocaine-conditioned mouse brain 1.5 h after post-test 2 on WD14 (Figure 6a). The number of c-Fos-positive cells—a marker of neuronal activity— was analyzed by immunohistochemistry. Compared with saline-conditioned mice, the number of c-Fos- and mVenus-double positive cells significantly increased in the NAc of cocaine-conditioned mice, indicating an increase in accumbal D1R-MSN activity (Figure 6b,c). The increase of c-Fos expression was suppressed by repeated ICA-27243 administration (Figure 6b,c).

**Figure 6.**
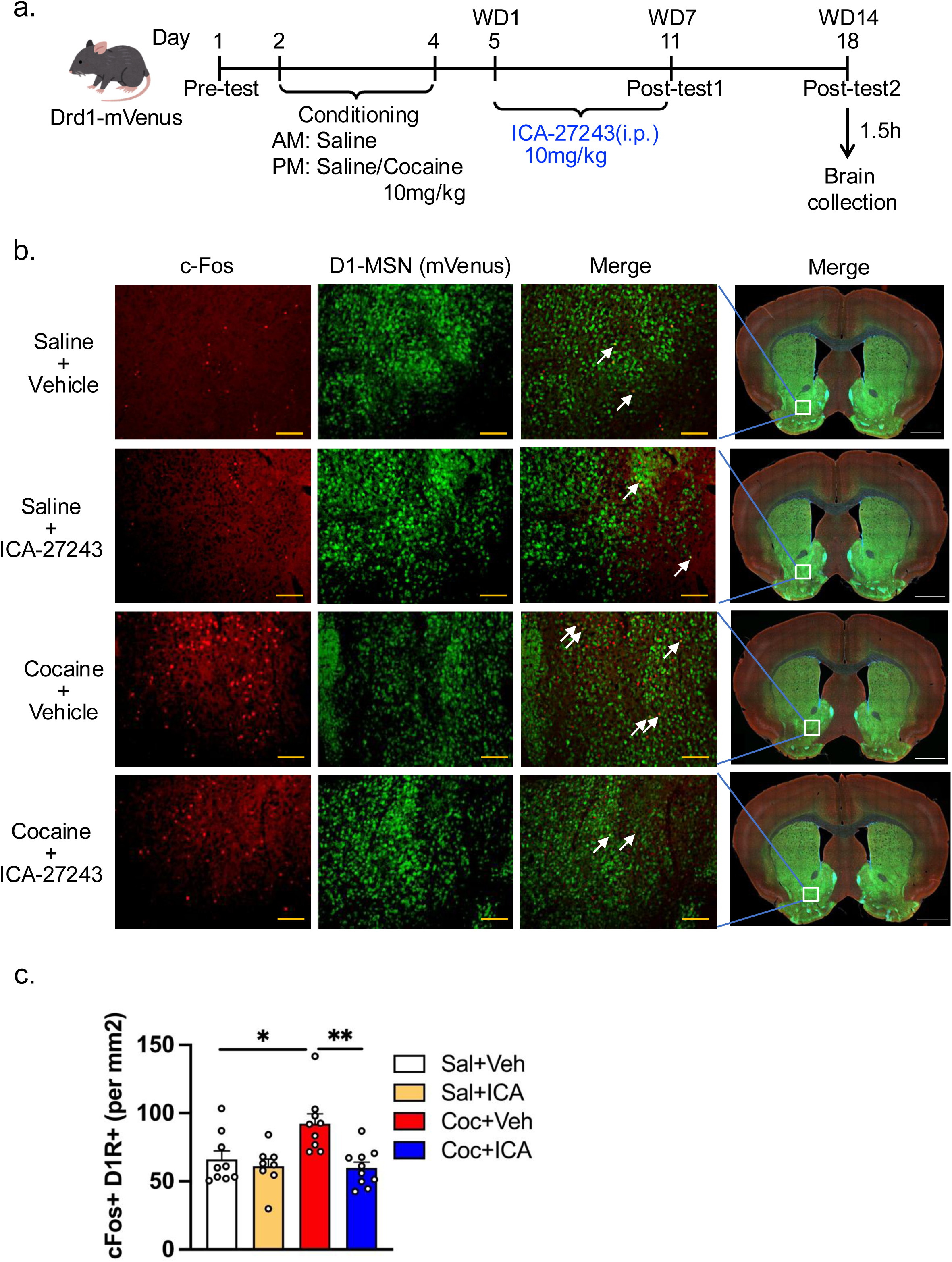
KCNQ2/3 potassium channel opener attenuates the cocaine-induced increase in the number of c-Fos-positive accumbal D1R-MSNs. **a.** Timeline of experiment procedure. D1-YFP mice were administered ICA-27243 or vehicle intraperitoneally for seven days, commencing on the first day of cocaine withdrawal (WD1). Post-tests were conducted on WD7 (day 11) and WD14 (day 18). Mouse brains were collected 1.5 h after post-test 2 on WD14. **b.** Representative fluorescent microscopic images showing c-Fos-positive cells in the NAc of D1-YFP mice. The yellow scale bar indicates 100 μm. The white scale bar indicates 1000 μm. **c.** Quantification of c-Fos-positive D1R-MSN cells per mm2. Two-way ANOVA: Cocaine, F(1,32)=10.4, P<0.01; ICA-27243, F(1,32)=4.508, P<0.05; interaction, F (1,32)=5.395, P<0.05. Tukey’s post-hoc test, *p<0.05, **p<0.01.

Next, we monitored the real-time activity of D1R-MSNs in mice during post-tests of the CPP test. We injected AAV-CAG-Flex-GCaMP6f into the NAc of Drd1-Cre mice and recorded the Ca^2+^ response in D1R-MSNs using fiber photometry during post-tests on WD7 and WD14 (Figure 7a). The Ca^2+^ response of D1R-MSNs increased just before mice entered the cocaine-paired box on WD7 and WD14 (Figure 7b−d), as previously reported^8^. ICA-27243 administration significantly attenuated the activity of D1R-MSNs in cocaine-conditioned mice on WD14, whereas it failed to attenuate the increased activity of D1R-MSNs on WD7 (Figure 7b−d). In saline-conditioned vehicle-treated mice, the activity of D1R-MSNs remained unchanged after mice entered the saline-paired box on WD7 and WD14 (Figure 7b−d). These findings suggest that KCNQ2/3 potassium channel opener inhibits abnormal accumbal D1R-MSN activity and cocaine-seeking behavior in mice.

**Figure 7.**
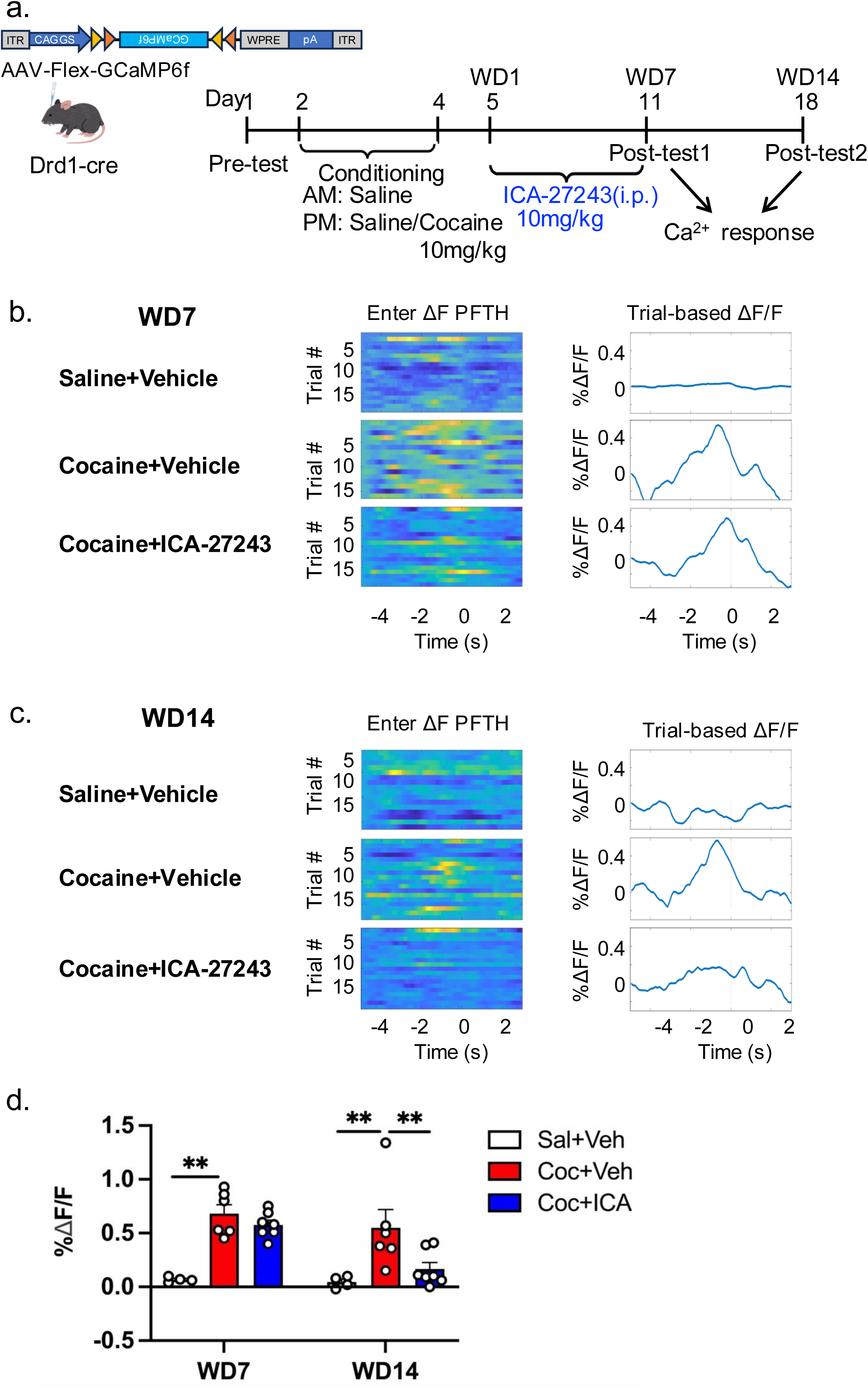
KCNQ2/3 potassium channel opener attenuates the cocaine-induced enhancement of D1R-MSN activity during cocaine-seeking behavior. **a.** Timeline of experimental procedure. Drd1-Cre mice were microinjected AAV-Flex-GCaMP6f into the NAc. The CPP test was carried out three weeks later. Mice were administered ICA-27243 or vehicle intraperitoneally for seven days, commencing on the first day of cocaine withdrawal (WD1). Post-tests were conducted on WD7 (day 11) and WD14 (day 18). During the post-test, the activity of D1R-MSNs was recorded by fiber photometry. **b,c.** Representative GCaMP6f signal in the accumbal D1R-MSNs on WD7 (b) and WD14 (c). Left panel—warmer colors on the heatmap represent higher fluorescent signals at that time point. Right panel—trial-based ΔF/F PETH before and after entering the cocaine-paired compartment. Time 0 indicates entry in the cocaine-paired compartment. **d.** Quantification of the peak amplitude of the Ca^2+^ signal in the 5 s preceding saline/cocaine-paired chamber entry. Two-way ANOVA: Drug effect, F(2, 14)=17.47, P<0.01; time effect, F(1, 14)=5.598, P<0.05; interaction effect, F(2, 14)=0.1423, P=0.1423; Tukey’s post-hoc test: **p<0.01. Saline+Vehicle group (n=4), Cocaine+Vehicle group (n=6), Cocaine+ICA-27243 group (n=7).

## DISCUSSION

Drug addiction is a chronic relapsing disorder that lacks a fundamentally effective pharmacological treatment^1,2^. Relapse during abstinence, driven by drug craving and seeking, remains a major therapeutic challenge, and no medication has hitherto succeeded in reliably preventing this transition^3^. Addressing this issue, our study strongly suggests that the KCNQ2/3 potassium channel is a master regulator of synaptic remodeling underlying cocaine-seeking behavior, as well as a promising therapeutic target for addiction.

We confirmed previous reports^9−13^ that the emergence of drug seeking after withdrawal (Figure 1b,d) coincides with a cluster of neuroadaptations in D1R-MSNs (Figure 3d), including increased thin spine formation (Figure S4a), enhanced surface expression of CP-AMPARs (Figure 4c−h), and heightened neuronal excitability (Figures 6 and 7). Interestingly, all of these behavioral, biochemical and morphological phenotypes were reversed by repeated administration of KCNQ2/3 channel openers or over-expression of a constitutively active KCNQ2 mutant (Figures 1 and 2), highlighting the central role of KCNQ2/3 in maintaining excitability and preventing maladaptive synaptic changes.

Previous studies have shown that repeated cocaine exposure promotes aberrant synaptogenesis in the NAc, including the formation of thin, silent synapses and the membrane insertion of CP-AMPARs^12,20,23^. These structural and functional plasticities are believed to support long-lasting drug-associated memories^7^. Our findings suggest that not only the induction but also the maintenance of such synaptic changes may depend on sustained or rhythmic neuronal firing in D1R-MSNs during the withdrawal period^15,27^. In this context, activation of KCNQ2/3 channels, which reduce membrane excitability and suppress firing frequency, may exert its therapeutic effect by disrupting the persistent electrical activity required to consolidate or maintain these maladaptive synaptic states^28^.

Importantly, the two KCNQ2/3 openers used in this study—ICA-27243 and XEN1101—were originally developed as antiepileptic drugs^29,30^. Their ability to dampen neuronal hyperexcitability is consistent with our proposed mechanism in addiction, where sustained D1R-MSN firing drives pathological synaptic remodeling^31^. Notably, phase 2 clinical trials for epilepsy have been completed for XEN1101, providing an established safety and pharmacokinetic profile in humans^30^. This significantly enhances its translational potential, as it could be repurposed for addiction treatment more rapidly than compounds at earlier stages of development^32^. ICA-27243, while less advanced clinically, further supports the mechanistic validity of targeting KCNQ2/3 to suppress relapse vulnerability^28^. Together, these pharmacological data not only underscore the feasibility of targeting KCNQ2/3 in humans but also reinforce the mechanistic conclusion drawn from our experiments.

Beyond cocaine, our findings may have broader implications for other addictive substances that elevate brain dopamine levels or induce sustained activation of NAc neurons^27,33^. Psychostimulants such as methamphetamine and amphetamine increase extracellular dopamine via transporter reversal or reuptake inhibition, leading to prolonged D1R-MSN activation^34,35^. Opioids, including heroin and prescription analgesics, indirectly enhance dopamine release by disinhibiting dopaminergic neurons in the ventral tegmental area^36,37^. Nicotine, acting through nicotinic acetylcholine receptors, and alcohol, via multiple neuromodulatory pathways, also promote persistent NAc neuronal activation^38,39^. In each of these cases, the resulting hyperdopaminergic state may trigger maladaptive synaptic remodeling similar to that observed with cocaine^17^.

Therefore, KCNQ2/3 channel activation could represent a generalizable strategy for reducing pathological excitability and relapse vulnerability across multiple forms of substance use disorders^18^.

Building on these clinical and preclinical observations, we propose that KCNQ2/3 serves as a master regulator of withdrawal-induced plasticity in D1R-MSNs. Structural plasticity in the form of de novo silent synapse formation and functional plasticity via CP-AMPAR signaling appear to arise from sustained KCNQ2/3 channel deactivation, likely driven by hyperdopaminergic tone^40,41^. While downstream dopamine receptor signaling pathways and metabotropic cascades may be contributing factors, our data suggest that failure to restore intrinsic excitability is a core pathophysiological driver of relapse vulnerability^42^.

Our findings also provide insight into the slow, progressive nature of the withdrawal state. Drug-seeking behavior and related synaptic abnormalities emerged only after 14 days of withdrawal (Figures 1−3), supporting a “threshold accumulation” model of relapse in which synaptic instability builds gradually before crossing a behavioral tipping point^12,23,43^. Whether this reflects transcriptional reprogramming or a progressive ionic/metabolic imbalance remains to be clarified.

Critically, our results reposition addiction as not only a disorder of synaptic plasticity but also of impaired membrane repolarization^42,44^. We propose that KCNQ2/3 channel dysfunction creates permissive conditions for synaptic remodeling, acting upstream of silent synapse maturation and CP-AMPAR recruitment (Figures 3−5). By suppressing spontaneous or stimulus-evoked firing that may be required to stabilize these synaptic changes, KCNQ2/3 activation could interrupt the positive feedback loop that sustains relapse vulnerability (Figures 1 and 2).

In conclusion, our study identifies KCNQ2/3 channels as a critical biological checkpoint in the pathogenesis of cocaine relapse. Given the lack of effective relapse prevention strategies^2^, these findings highlight the therapeutic potential of KCNQ2/3 channel openers—particularly XEN1101, with its completed Phase 2 program^30^—in providing a realistic and accelerated path toward clinical application. By restoring membrane stability, reducing pathological firing, and suppressing aberrant synaptic plasticity (Figure 8), modulation of KCNQ2/3 may offer a novel, translatable route to pharmacological intervention for substance use disorders, which remains an urgent global challenge.

**Figure 8.**
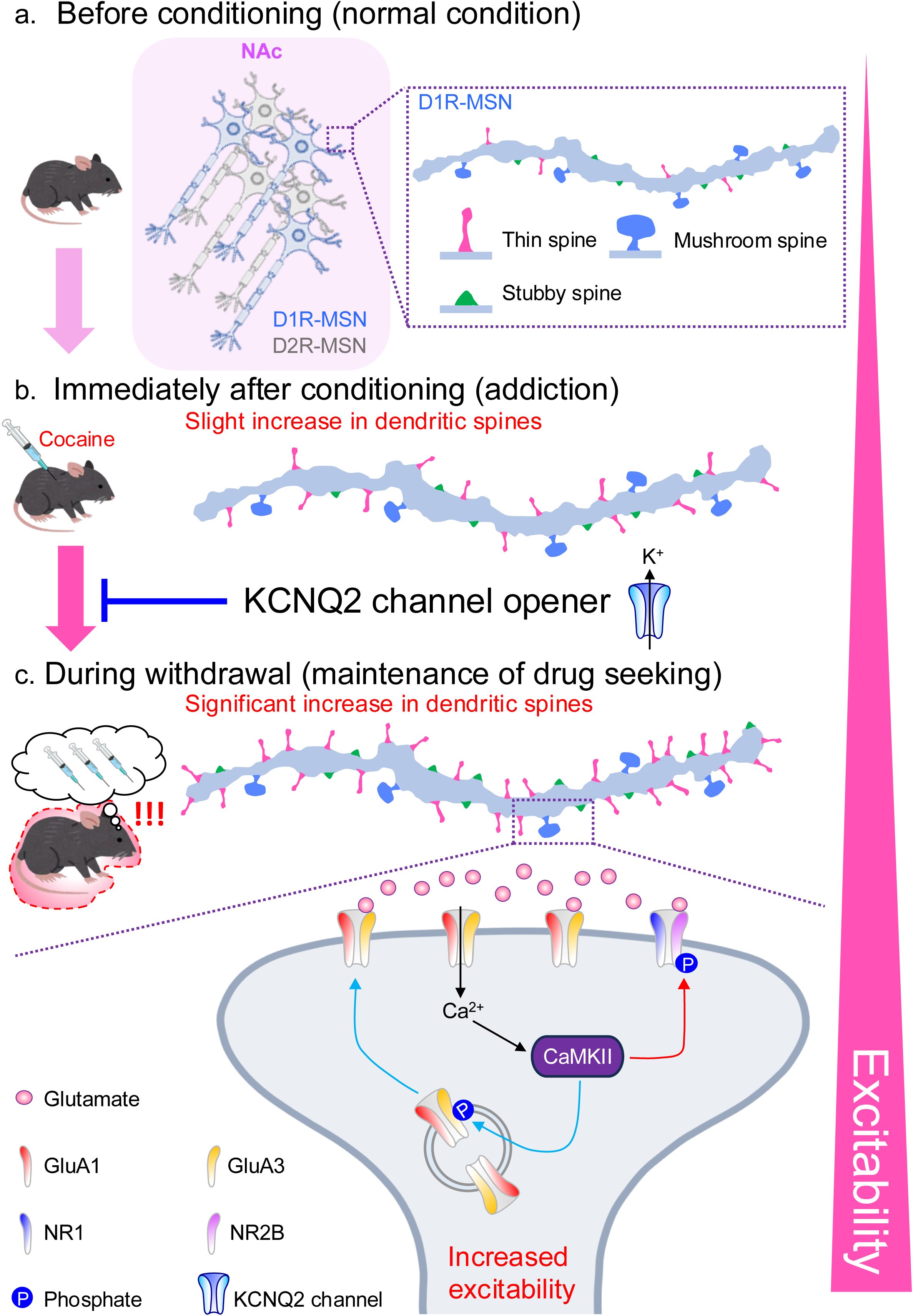
Working model of the ameliorating effect of a KCNQ2/3 channel opener on drug-seeking behavior during cocaine withdrawal. **a.** Before conditioning—spine density is at basal levels. **b.** Immediately after conditioning—the number of dendritic spines slightly increases, with minimal CP-AMPAR expression. **c.** During withdrawal—both the number of dendritic spines (particularly thin spines) and cell surface CP-AMPAR expression increase significantly, and neuronal excitability also increases. When glutamate stimulation activates CP-AMPAR, Ca²⁺ flows into the neuron, in turn activating CaMKII. This promotes the phosphorylation of GluA1 (S831) and NR2B (S1303), thereby enhancing synaptic plasticity and contributing to the maintenance of drug-seeking behavior. Administration of KCNQ2 channel openers to mice during withdrawal effectively inhibits neuronal excitability, suppresses cell surface CP-AMPAR upregulation and synaptic plasticity, and ultimately reduces drug-seeking behavior.

Although the results are encouraging, there are several limitations to this study. First, while both behavioral and molecular findings strongly support the role of the KCNQ2 potassium channel in regulating cocaine-seeking behavior during withdrawal in mice, the specific downstream signaling pathways remain unclear. Second, this study only used the CPP addiction model; thus, generalizability of these findings to other addiction and withdrawal models requires further investigation. Additionally, given that KCNQ2 channels are widely distributed in the central nervous system, long-term use of openers may impact normal neuronal activity, including for learning and memory. Although no significant cognitive impairment has been observed so far, further studies using tests such as the water maze or novel object recognition are needed to assess long-term safety.

## METHODS

### Experimental model and sample details

#### Animals

Eight-week-old male C57BL/6 mice (weighing 20−25 g) were purchased from Japan SLC, Inc. (Shizuoka, Japan). Drd1-Cre and Adora2a-cre mice on a C57BL/6 background were obtained from the Mutant Mouse Research and

Resource Center (MMRRC) at the University of California, Davis (Davis, CA, USA). Transgenic mice expressing a variant of yellow fluorescent protein (mVenus) under the control of the D1R promoter (Drd1-mVenus, RBRC03111, Riken BRC, Tsukuba, Japan) were genotyped by polymerase chain reaction (PCR) using genomic DNA extracted from tail clips. The mice were randomly assigned to groups of four per cage and housed under standard conditions: a 12-h light/dark cycle (lights on from 08:00 to 20:00), a constant temperature of 23 ± 1°C, and humidity maintained at 50 ± 5%. Food and water were available ad libitum throughout the experiments. All experiments were conducted during the light phase.

### Procedure

#### CPP test

The CPP test was performed as previously described, with minor modifications^14^. The CPP apparatus consisted of three compartments. One compartment featured a black-and-white striped wall with a smooth surface floor (width 24 cm × depth 33 cm × height 18 cm), while the other compartment had a white wall with a textured floor (width 24 cm × depth 33 cm × height 18 cm). A small transitional compartment (width 12 cm × depth 33 cm × height 18 cm) connected the two main compartments, with doors that could be controlled to regulate access.

The CPP test comprised three distinct phases: pre-test, conditioning phase, and post-test. During the pre-test, mice were placed in the apparatus and allowed to freely explore all three compartments for 15 min. The time spent in each compartment was recorded automatically (Med Associates Inc, Vermont, USA).

Mice that spent more than 600 s in a single compartment were excluded from further analysis due to location bias.

During the conditioning phase, each mouse was randomly assigned to one of two compartments—either the black-and-white striped compartment or the white compartment—and confined there for 30 min immediately after receiving a saline injection in the morning. In the afternoon, the same mouse was confined to the opposite compartment for 30 min following the administration of either saline or cocaine.

During the post-test, mice were again placed in the apparatus and allowed to explore all three compartments freely for 15 min, with the time spent in each compartment recorded. The CPP score was calculated by subtracting the time spent in the cocaine-paired compartment during the pre-test from the time spent in the same compartment during the post-test.

Mice were administered ICA-27243, XEN1101, ICA-110381, or a combination of XE991 and ICA-27243 between the day following the conditioning phase and the first post-test. Additionally, Naspm was administered prior to the second post-test.

#### Stereotaxic surgery

The mice were intraperitoneally administered an anesthetic mixture consisting of 0.75 mg/kg medetomidine (Nippon Zenyaku Kogyo, Fukushima, Japan), 4 mg/kg midazolam (Sandoz, Basel, Switzerland), and 5 mg/kg butorphanol (Meiji Seika Pharma, Tokyo, Japan). Once fully anesthetized, each mouse was secured in a stereotaxic apparatus. A midline incision was made along the superior aspect of the cranium to expose the skull. A total volume of 0.5 μl of AAV solution (1.0×10¹² genome copies/mL) was bilaterally microinjected into the NAc at a rate of 0.1 μl/min (two injection sites per side) through a hole drilled into the skull. The injection coordinates were as follows: AP +1.6 mm, ML ±1.5 mm, DV −4.4 mm; and AP +1.0 mm, ML ±1.0 mm, DV −4.5 mm from the bregma, with an injection angle of 10°.

For Naspm microinjection, mice were anesthetized and stereotaxically implanted with a guide and dummy cannula (Eicom, Kyoto, Japan) targeting the NAc (bregma coordinates: AP +1.3 mm, ML ±1.5 mm, DV −3.5 mm) 24 h after post-test 1. Six days after surgery, the dummy cannula was replaced with a microinjection cannula (1 mm longer than the guide cannula), and 40 μg of Naspm [dissolved in 0.5 μl phosphate-buffered saline (PBS)] was microinjected into the NAc over a 5-min period. Post-test 2 was conducted 10 min after the microinjection.

#### Immunohistochemistry

The mice were anesthetized and perfused transcardially with ice-cold PBS followed by 4% paraformaldehyde (PFA). The brains were extracted and post-fixed in 4% PFA overnight at 4°C. Subsequently, the brains were transferred to a 30% sucrose solution for cryoprotection. Coronal brain sections (30 μm-thick) containing the NAc were obtained using a sliding microtome (Leica, Wetzlar, Germany). The sections were fixed in 4% PFA for 5 min, then permeabilized with PBS containing 0.3% Triton X-100 for 10 min. After two 10-min washes in PBS, the sections were blocked for 1 h in a blocking solution containing 5% normal donkey serum. Following three additional 10-min PBS washes, the sections were incubated overnight at 4°C with a rabbit anti-GFP antibody (1:2000). The next day, sections were washed three times (5 min each) in PBS and incubated at room temperature for 1 h with a Goat Alexa Fluor 488-labeled anti-rabbit IgG antibody (1:1000). After three final 10-min PBS washes, the sections were mounted on glass slides.

For c-Fos immunofluorescence staining, the sections were incubated overnight at 4°C with a rabbit anti-c-Fos antibody (1:1000). The next day, they were washed three times (5 min each) in PBS and incubated at room temperature for 1 h with either a Goat Alexa Fluor 594-labeled anti-rabbit IgG antibody (1:1000) or a Goat Alexa Fluor 488-labeled anti-rabbit IgG antibody (1:1000).

#### Spine analysis

Mice that had been microinjected with AAV-Sp-Cre and AAV-Flex-EGFP into the NAc were transcardially perfused with 4% PFA immediately after the CPP test. The brains were sectioned into 150-μm-thick slices using a vibratome (Leica). Subsequently, the sections were immunostained with a rat anti-GFP antibody, followed by a Goat Alexa Fluor 488-labeled anti-rat IgG antibody. Images were acquired using a laser confocal microscope (LSM 980, Zeiss, Oberkochen, Germany) with a 63× oil immersion objective and a frame size of 512 × 512 pixels. Accumulated secondary dendrites of MSNs were selected from branches located at least 50 μm away from the cell body. For each neuron, three secondary dendritic spines were analyzed on 2–3 neurites, covering approximately 150 μm of secondary dendrite. We examined three neurons per mouse, with a total of six mice per group.

Dendritic spines were classified into three categories^45^: mushroom spines (maximum width > 0.6 μm); stubby spines (maximum width < 0.6 μm and length/maximum width < 1); and thin spines (maximum width < 0.6 μm, length/maximum width > 1, and length < 2 μm). Spine density for each subtype, as well as the total spine density, was analyzed using Imaris software (Bitplane, Belfast, United Kingdom).

#### Preparation of striatal slices

Striatal slices were prepared as described previously^46^. The brains were removed and stored in cold Krebs-HCO3 buffer (124 mM NaCl, 4 mM KCl, 26 mM NaHCO3, 1.5 mM CaCl2, 1.25 mM KH2PO4, and 10 mM D-glucose, pH 7.4). Coronal brain slices (350 µm) were prepared using a vibratome. The entire striatum and NAc were then dissected from the slices and incubated in Krebs-HCO3 buffer containing adenosine deaminase (10 µg/ml) for 30 min. The slices were subsequently treated with (S)-AMPA (10 µM) for 15 s. After treatment, the slices were flash-frozen in liquid nitrogen and stored at −80°C until immunoblotting experiments were performed.

#### In vivo TurboID protein purification

In vivo TurboID experiments were performed as previously described^47^, with minor modifications. AAV-Ef1a-DIO-TurboID-surface was bilaterally microinjected into the NAc. Three weeks after viral injection, mice underwent the CPP behavioral test. Following the post-test, biotin was subcutaneously administered at a dose of 50 mg/kg. The NAc was collected 1 h after biotin administration. Each NAc was lysed in a buffer containing 50 mM Tris/HCl (pH 7.5), 150 mM NaCl, 1 mM EDTA, 0.2% SDS, 1% TritonX-100, 1% deoxycholate, protease inhibitor mixture (cOmplete Mini EDTA-free, Roche), and phosphatase inhibitor mixture (PhosSTOP, Roche). The lysate was then sonicated and centrifuged at 15,000g for 10 min. The cleared supernatant was supplemented with SDS to a final concentration of 1% and heated at 55°C for 15 min. After cooling on ice, the sample was incubated overnight at 4°C with Pierce High Capacity NeutrAvidin Agarose (Thermo Fisher). Beads were sequentially washed twice with 2% SDS, twice with 1% Triton X-100 and 1% deoxycholate in 25 mM LiCl, twice with 1 M NaCl, and five times with 50 mM ammonium bicarbonate. Biotinylated proteins were eluted in 125 mM Tris-HCl (pH 6.8), 4% SDS, 0.2% β-mercaptoethanol, 20% glycerol, and 3 mM biotin at 60°C for 15 min.

#### Immunoblotting

The samples (10 μg) were loaded onto an 8% SDS-PAGE gel and subjected to electrophoresis. The proteins were transferred onto a polyvinylidene fluoride membrane (Merck, Darmstadt, Germany). The membrane was then blocked with Blocking One (Nacalai Tesque Inc., Kyoto, Japan) at room temperature for 1 h. Following blocking, the membrane was washed three times with Tris-buffered saline containing 0.1% Tween 20 (TBST) for 10 min per wash. It was then incubated overnight at 4°C with primary antibodies: rabbit anti-phospho-GluA1 (S831) (1:1000), mouse anti-GluA1 (1:1000), rabbit anti-phospho-NR2B (S1303) (1:1000) and mouse anti-NR2B (1:1000), rabbit anti-GluA2 (1:1000) and rabbit anti-GluA3 (1:1000). After incubation, the membrane was washed again three times with TBST for 10 min each. Subsequently, the membrane was incubated at room temperature for 1 h with secondary antibodies: Goat Alexa Fluor 680-labeled anti-rabbit IgG and Goat DyLight 800-labeled anti-mouse IgG (H+L). The membrane was washed three more times with TBST for 10 min each. Finally, protein bands were visualized and analyzed using the Odyssey CLx imaging system (LI-COR Inc., Lincoln, USA).

#### Fiber photometry Ca^2+^ imaging

pAAV.CAG.Flex.GCaMP6f.WPRE.SV40 was a gift from Douglas Kim & GENIE Project (Addgene viral prep #100835-AAV9; http://n2t.net/addgene:100835; RRID:Addgene_100835). AAV-CAG-Flex-GCaMP6f^48^ was microinjected into the NAc of Drd1-Cre mice at the following stereotaxic coordinates: AP +1.4 mm; ML +1.5 mm; DV −4.4 mm (10° angle, vertical). A fiber-optic cannula was implanted just above the viral injection site (AP +1.4 mm; ML +1.5 mm; DV −4.3 mm; 10° angle) and secured to the skull using a light-cured bonding adhesive and composite resin (Kuraray Noritake Dental Inc., Japan). Mice were allowed to recover for three weeks before behavioral handling. In vivo Ca²⁺ imaging was conducted using the TeleFipho wireless fiber photometry system (Bioresearch Center Inc., Aichi, Japan). Behavioral events, including paired and unpaired chamber entries, were recorded in real time via transistor−transistor logic (TTL) signals generated by Med-PC Behavioral Control software. For each mouse, the time of entry into the drug-paired chamber was identified, and fluorescence signals within a 5-s window surrounding the entry were analyzed^8^. Changes in fluorescence (ΔF/F) were processed using the open source software PMAT^49^.

### Quantification and statistical analysis

All data are presented as mean ± standard error of the mean (SEM). Statistical analyses were performed using GraphPad Prism 9 (GraphPad Software, Boston, USA). One-way or two-way ANOVA was conducted, followed by Tukey’s post-hoc test for multiple comparisons when a significant main effect was detected (p<0.05).

## RESOURCE AVAILABILITY

### Lead contact

Requests for further information and resources should be directed to and be fulfilled by the lead contact, Taku Nagai.

### Materials availability

This study did not generate new unique reagents.

### Data and code availability

All data have been deposited at Mendeley Data and are publicly available as of the date of publication at [DOI: 10.17632/x3jt94d2pm.2].

This paper does not report original code.

Any additional information required to reanalyze the data reported in this paper is available from the lead contact upon request.

## ACKNOWLEDGMENTS

We thank all members of the Nagai and Kaibuchi laboratories for their technical support and valuable discussions. We are sincerely grateful to Professor Hisatsugu Koshimizu of the Office of Research Administration at Fujita Health University for the valuable support in refining our manuscript. We also thank the Open Facility Center and Advanced Medical Research Center of Animal Models for Human Diseases at Fujita Health University.

This work was supported by the following funding sources: AMED Grant Number [JP21wm0425017, T. Nagai (T. N.); JP21wm0425008, T. N.; JP23tm0524001, T. N.]; JSPS KAKENHI Grant Number [23H02843, T. Nabeshima; 23K27360, K. Y.; 24K02218, T. N.; 24K18291, X. Zhang; 25K09861, Y. F.]; JSPS J-PEAKS (K. K.; T. N.); GRANT-IN-AID from FHU (T. N.; X. Zhang); CSC [Grant No. 202308050112, X. Zhou]; JST PRESTO [21461219, T. T.; 24029397, T. T.]; AMED Brain/MINDS 2.0 [24019272, T. T; 24019528, T. T.]; the Ono Pharmaceutical Foundation [Oncology, Immunology and Neurology, T. T.]; the Memorial Foundation for Medical and Pharmaceutical Research (T. T.); the JGC Saneyoshi Scholarship Foundation (T. T.); the Takeda Science Foundation (T. T.); and MEXT Grants-in-Aid for Scientific Research on Innovative Areas “Cell type census of adaptive neuronal circuits: biological mechanisms of structural and functional organization (Adaptive Circuit Census, ACC)” [24H01250, T. T.].

## AUTHOR CONTRIBUTIONS

X. Zhou: Writing – original draft, writing – review & editing, conceptualization, data curation, research data, investigation, methodology, visualization, validation. X. Zhang: Writing – original draft, writing – review & editing, conceptualization, data curation, research data, investigation, methodology, resources, visualization, validation. Y. F.: Investigation, resources. D.T.: Methodology, resources. T. T.: Methodology, resources. H. K.: Resources. C. T. Y.: Writing – review & editing. T. Nabeshima: Review, resources. K. Y.: Methodology. K. K.: Resources, project administration, supervision. T. Nagai: Writing – review & editing, conceptualization, project administration, supervision, visualization, funding acquisition.

## DECLARATION OF INTERESTS

All authors declare no competing interests for this work.

## DECLARATION OF GENERATIVE AI AND AI-ASSISTED TECHNOLOGIES

During the preparation of this work, the authors used ChatGPT to improve language. After using this tool or service, the authors reviewed and edited the content as needed and take full responsibility for the content of the publication.

## EXTENDED DATA

### STAR★Methods

#### Key resources table

| REAGENT OR RESOURCE | SOURCE | IDENTIFIER |
| --- | --- | --- |
| Antibodies |  |  |
| Anti-GFP (Rat IgG2a), Monoclonal (GF090R) | Nacalai Tesque | Cat# 04404-26, RRID: AB_10013361 |
| Anti-c-Fos | Synaptic Systems | Cat# 226003, RRID: AB_2231974 |
| Rabbit monoclonal anti-KCNQ2 (D9L5S) antibody | Cell Signaling Technology | Cat# 14752, RRID: AB_2798597 |
| F(ab')<sub>2</sub>-Goat anti-Rabbit IgG (H+L) Cross-Adsorbed<br>Secondary Antibody, Alexa Fluor™ 594 | Thermo Fisher Scientific | Cat# A-11072, RRID: AB_2534116 |
| Goat anti-Rat IgG (H+L) Cross-Adsorbed Secondary<br>Antibody, Alexa Fluor™ 488 | Thermo Fisher Scientific | Cat# A-11006, RRID: AB_2534074 |
| Phospho-AMPA Receptor 1 (GluA1) (Ser831) (A5O2P)<br>Rabbit mAb | Cell signaling Technology | Cat# 75574, RRID: AB_2799873 |
| AMPA Receptor 1 (GluA1) (D4N9V) | Cell signaling Technology | Cat# 13185, RRID: AB_2732897 |
| AMPA Receptor (GluR2) (E1L8U) Rabbit mAb | Cell signaling Technology | Cat# 13607, RRID: AB_2650557 |
| AMPA Receptor (GluR3) (D25G9) Rabbit mAb | Cell signaling Technology | Cat# 5117, RRID: AB_10544796 |
| Goat anti-Rabbit IgG (H+L) Highly Cross-Adsorbed<br>Secondary Antibody, Alexa Fluor™ 680 | Thermo Fisher Scientific | Cat# A-21109, RRID: AB_2535758 |
| DyLight 800-labeled anti-mouse IgG(H+L) | Cell signaling Technology | Cat# 5257, RRID: AB_10693543 |
| Anti β-Actin | FUJIFILM Wako Pure Chemical<br>Corporation | Cat# 010-27841, RRID: AB_2858279 |
| Anti-RFP mAb | MBL International | Cat# M155-3, RRID: AB_1278880 |
| Rabbit monoclonal anti-phospho-NMDAR2B (Ser1303) | Abcam | Cat# ab81271, RRID: AB_1640731 |
| Mouse monoclonal anti-NMDAR2B | BD Biosciences | Cat# 610417, RRID: AB_397797 |
| Mouse monoclonal anti-GluR1-NT | Sigma-Aldrich | Cat# MAB2263, RRID: AB_11212678 |
| Goat anti-Rabbit IgG (H+L) Highly Cross-Adsorbed<br>Secondary Antibody, Alexa Fluor™ 555 | Thermo Fisher Scientific | Cat# A-21429, RRID: AB_2535850 |
| Bacterial and virus strains |  |  |
| AAV-Sp-Cre | Nagai et al. 2016 | N/A |
| AAV-CAGGS-Flex-EGFP | Nagai et al. 2016 | N/A |
| AAV-CAGGS-Flex-mCherry | Funahashi et al. 2019 | N/A |
| AAV-Ef1a-DIO-TurboID-HA-GPI | Takano et al. 2020 | N/A |
| AAV-CW3SL-Flex-KCNQ2-CA--mCherry | This study | N/A |
| pAAV-CAG-Flex-GCaMP6f-WPRE-SV40 | Douglas Kim & GENIE Project | RRID: Addgene viral prep # 100835-AAV9 |
| Chemicals, peptides, and recombinant proteins |  |  |
| Cocaine | Mallinckrodt | NDC# 0406-1520-55 |
| ICA-27243 | MedChemExpress | Cat# HY-122114 |
| ICA-110381 | Alomone Labs | Cat# I-160 |
| XE991 | Tocris Bioscienc | Cat# 2000 |
| Naspm | Tocris Bioscienc | Cat# 2766/10 |
| Kainic acid | Biomol GmbH | Cat# EA-123 |
| XEN1101 | MedChemExpress | Cat# HY-139791 |
| Experimental models: Organisms/strains |  |  |
| C57BL/6NcrSlc male | Japan SLC | RRID : MGI : 5295404 |
| Drd1-mVenus (C57BL/6J-Tg (mDrd1-YFP) 680-1Koba) | Nagai et al. 2016 | RRID: IMSR_RBRC03111 |
| Software and algorithms |  |  |
| GraphPad Prism 9 | GraphPad | https://www.graphpad.com/ |
| Med-PC Behavioral Control software | Med Associates, Inc. | https://med-associates.com/ |
| Photometry Modular Analysis Tool | Barker Lab, Rutgers University. | https://github.com/djamesbarker/pMAT |

---

**Figure S1.**
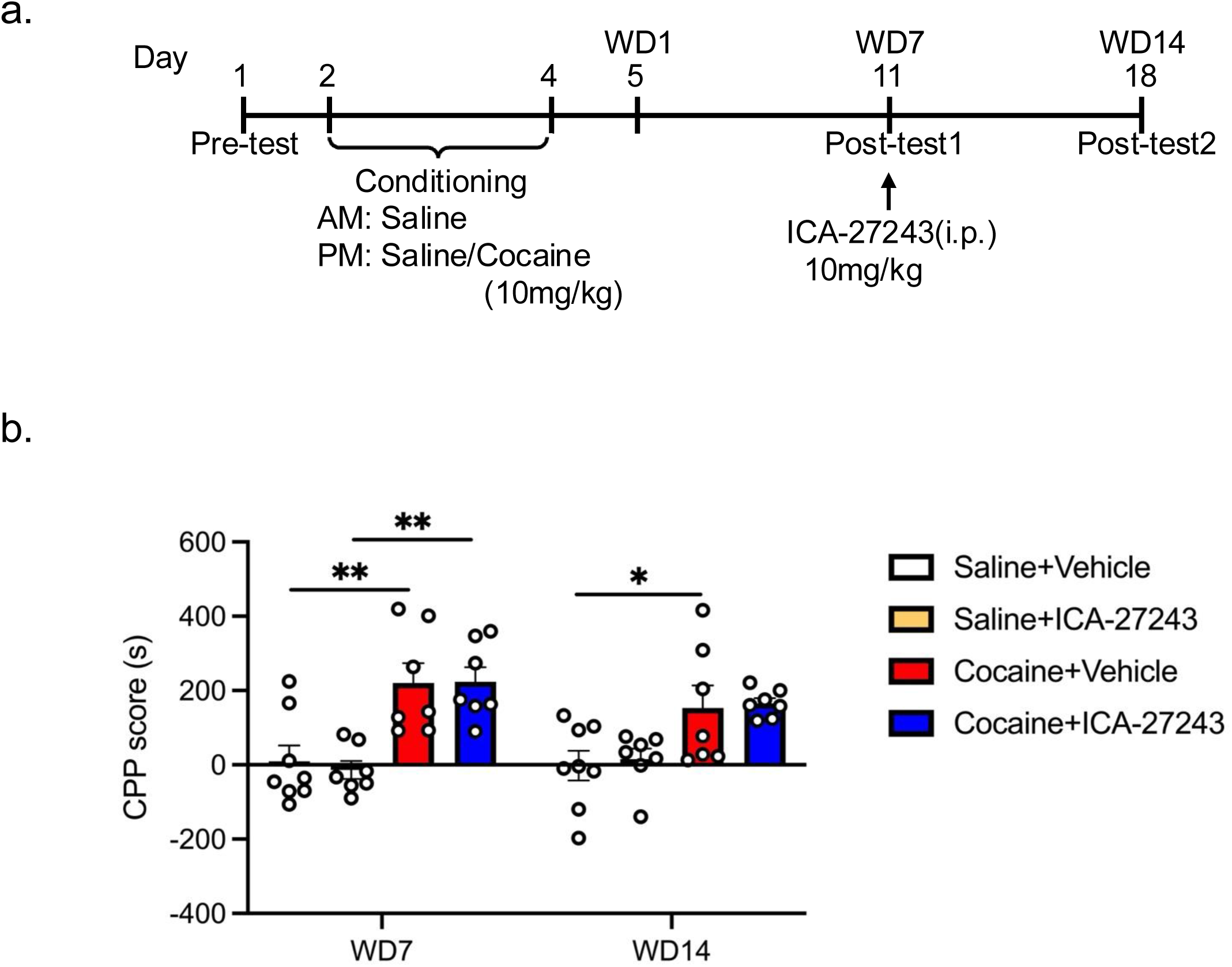
KCNQ2/3 potassium channel opener single treatment has no effect on cocaine-seeking behavior after 14-day withdrawal a,b. Effect of single ICA-27243 on cocaine-seeking behavior after 14-day withdrawal. **a.** Timeline of CPP procedure. Mice were administered ICA-27243 or vehicle 30min before post-test on WD7 (day11), commencing on the first day of cocaine withdrawal (WD1). Post-tests were conducted on WD7 (day11) and WD14 (day 18). **b.** CPP score of WD7 and WD14. The administration of ICA-27243 (10 mg/kg) had no effect on CPP scores. Two-way ANOVA: Drug effect, F(3, 25)=8.230, P=0.0006; time effect, F(1, 25)=3.294, P=0.0816; interaction effect, F(3, 37)=2.208, P=0.1121. Tukey’s post-hoc test: \**p*<0.05, \*\**p*<0.01. Saline+Vehicle group (n=8), Saline+ICA-27243 group (n=7), Cocaine+Vehicle group (n=8), Cocaine+ICA-27243 group (n=8).

**Figure S2.**
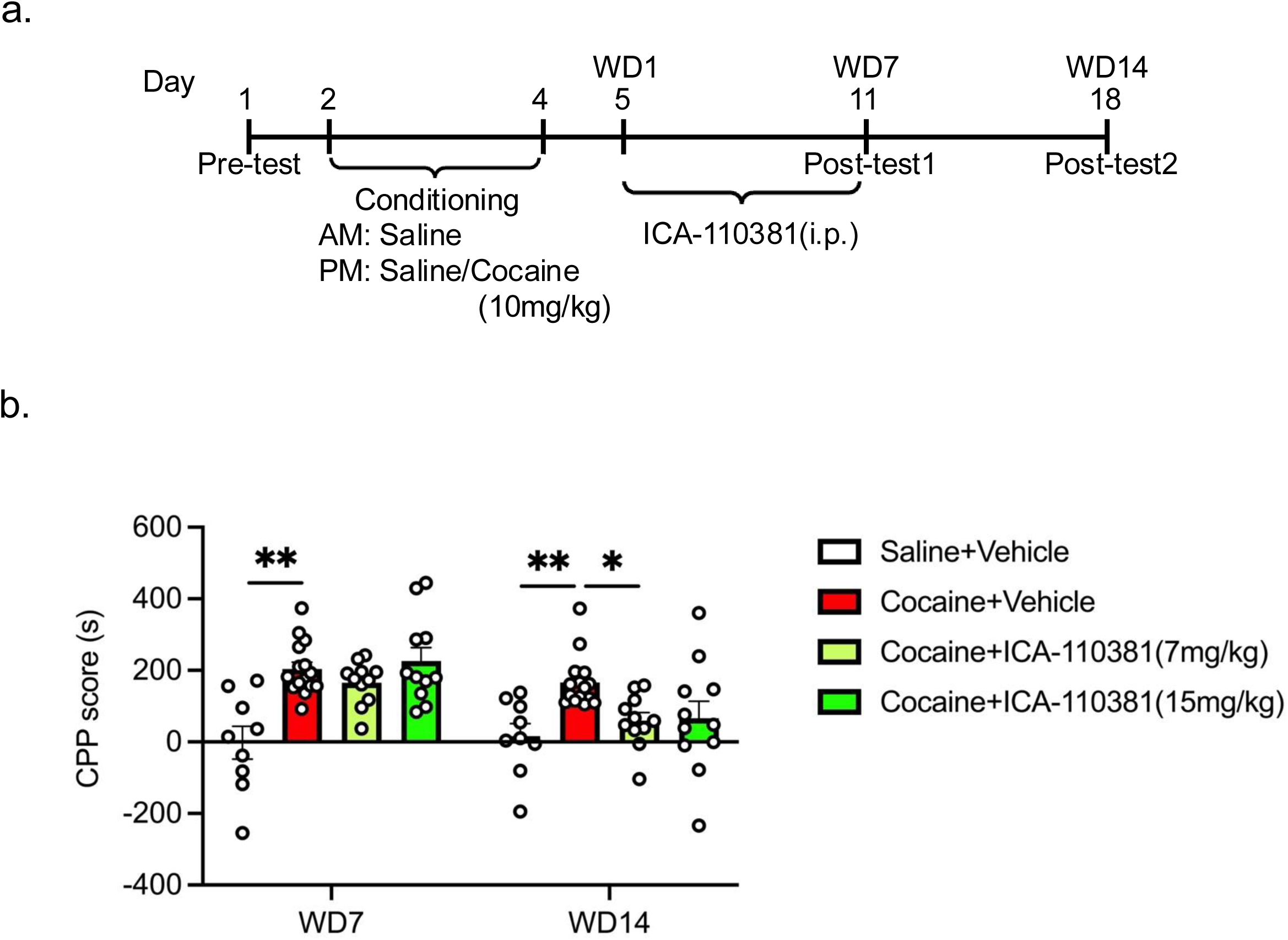
KCNQ2/3 potassium channel opener ICA-110381 attenuates cocaine-seeking behavior after 14-day withdrawal a,b. Effect of ICA-110381 on cocaine-induced CPP after 14-day withdrawal. **a.** Timeline of CPP procedure. Mice were administered ICA-110381 or vehicle intraperitoneally for seven days, commencing on the first day of cocaine withdrawal (WD1). Post-tests were conducted on WD7 (day 11) and WD14 (day 18). **b.** CPP score of WD7 and WD14. The administration of ICA-110381 (7 mg/kg) resulted in the attenuation of the CPP score induced by cocaine on WD14. Two-way ANOVA: Drug effect, F(2, 32)=15.17, P<0.001; time effect, F(1, 32)=8.957, P<0.01; interaction effect, F(2, 32)=5.978, P<0.01; Tukey’s post-hoc test: **p<0.01. Saline+Vehicle group (n=9), Cocaine+Vehicle group (n=15), Cocaine+ICA-110381(7mg/kg) group (n=11), Cocaine+ICA-110381(15mg/kg) group (n=11).

**Figure S3.**
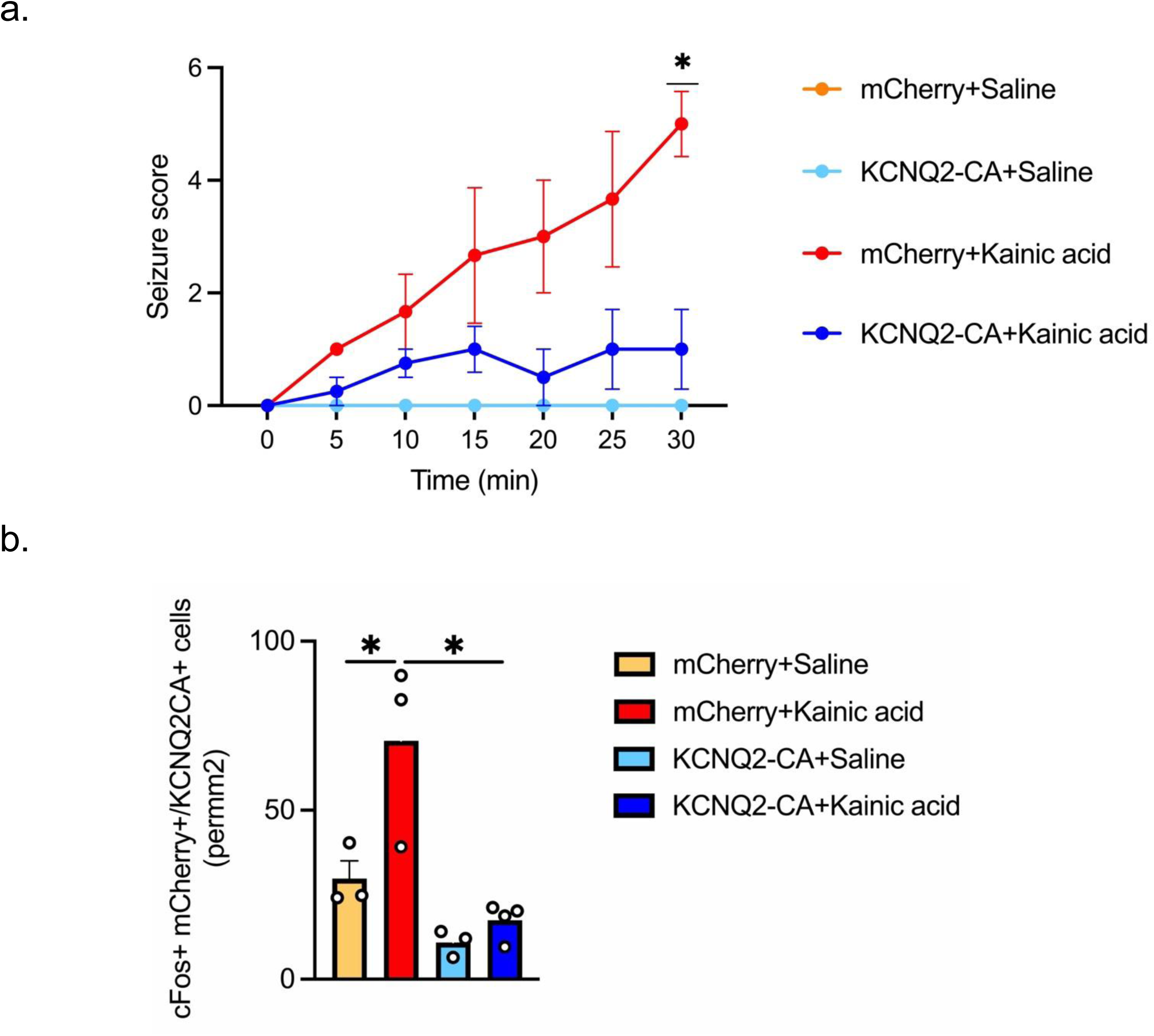
AAV-Flex-KCNQ2-CA significantly inhibits the excitability of D1R-MSNs in the NAc. **a.** After kainic acid (25mg/kg) injection, the mean behavioral seizure scores of AAV-Flex-KCNQ2-CA mice were significantly reduced. Three-way ANOVA: Time, F(1.87,20.57)=14.65, P=0.0001; kainic acid, F(1,11)=22.90, P=0.0006; KCNQ2-CA, F(1,11)=7.739 P=0.0178; time x kainic acid, F(6,66)=14.65, P<0.0001; time x KCNQ2-CA, F(6,66)=6.618, P<0.0001; kainic acid x KCNQ2-CA, F(1,11)=7.739 P=0.0178; time x kainic acid x KCNQ2-CA, F(6,66)=6.618, P<0.0001. **b.** Quantification of c-Fos-positive D1R-MSN per mm^2^. Two-way ANOVA: Kainic acid, F(1,8)=7.802, P=0.0234; KCNQ2-CA, F(1,8)=16.88, P=0.0034; interaction, F(1,8)=3.705, P=0.0904. Tukey’s post-hoc test, \**p*< 0.05.

**Figure S4.**
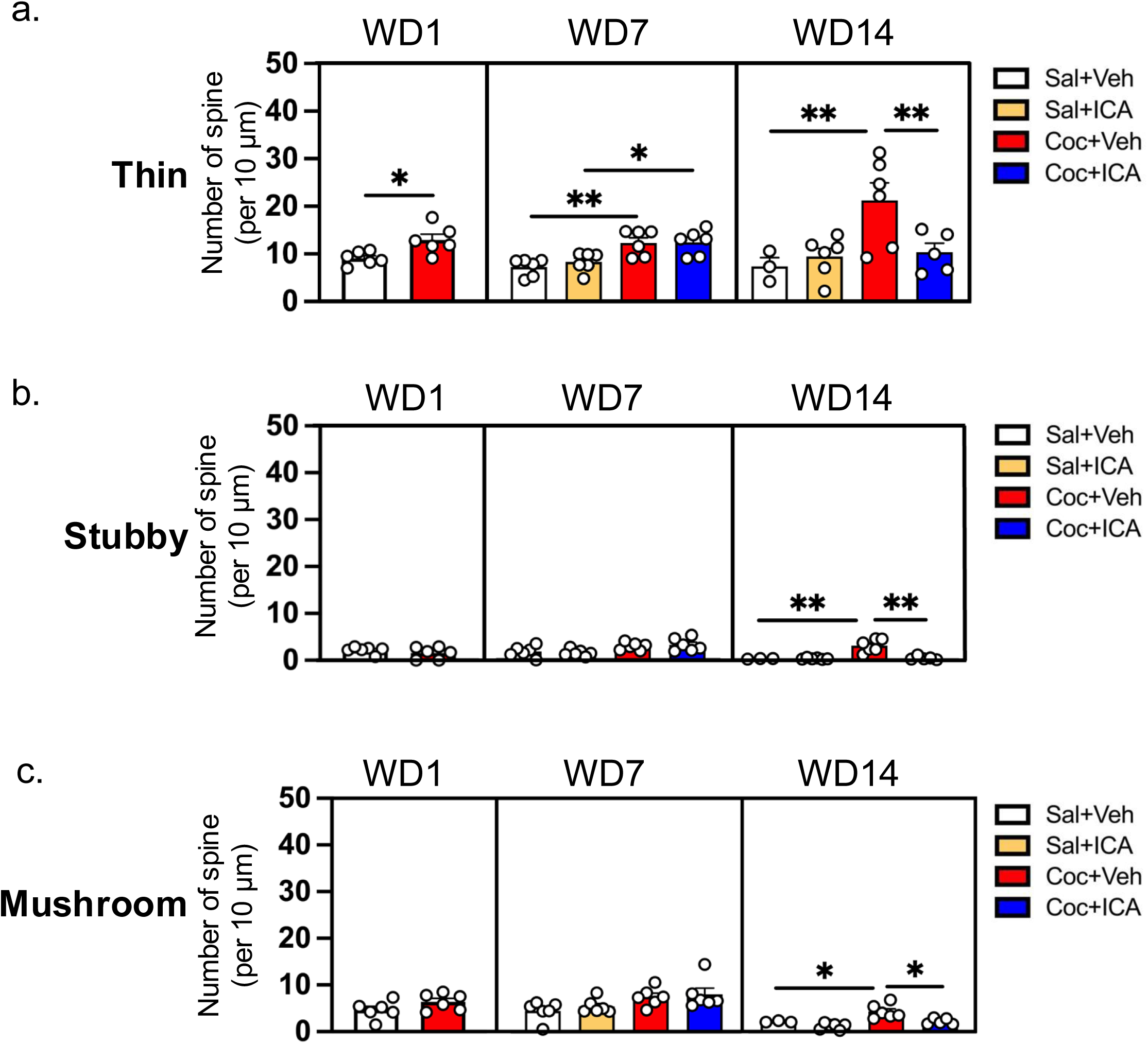
KCNQ2/3 potassium channel opener treatment attenuated the increase of spine density of thin and stubby induced by cocaine a−c. The quantification of the spine density on WD1, WD7, and WD14. **a.** Thin spine. Two-tailed unpaired t-test: WD1, t=2.923, P=0.0152. Two-way ANOVA: WD7, ICA-27243: F(1,20)=0.3764, P=0.5464; cocaine: F(1,20)=23.44, P<0.0001; interaction: F(1,20)=0.2526, P=0.6207; WD14, ICA-27243: F(1,20)=17.58, P<0.01; cocaine, F(1,20)=16.32, P<0.01; interaction, F(1,20)=8.685, P<0.01. Tukey’s post-hoc test; \**p*<0.05, \*\**p*<0.01. **b.** Stubby spine. Two-tailed unpaired t-test: WD1, t=1.092, P=0.3005. Two-way ANOVA: WD7, ICA-27243, F(1,20)=0.006706, P=0.9356; cocaine, F(1,20)=10.66, P<0.05; interaction, F(1,20)=0.4663, P=0.5025; WD14, ICA-27243, F(1,20)=10.44, P<0.01; cocaine, F(1,20)=8.61, P<0.01; interaction, F(1,20)=10.09, P<0.01. Tukey’s post-hoc test: **p<0.01. **c.** Mushroom spine. Two-tailed unpaired t-test: WD1, t=1.622, P=0.1358. Two-way ANOVA: WD7, ICA-27243, F(1,20)=0.6261, P=0.4381; cocaine, F(1,20)=8.590, P<0.01; interaction, F(1,20)=0.0459, P=0.8325; WD14, ICA-27243, F(1,20)=0.3702, P=0.5497; cocaine, F(1,20)=0.4393, P=0.515; interaction: F(1,20)=0.303, P=0.5881. Tukey’s post-hoc test: *p<0.05.

## Notes

### Competing Interest Statement

The authors have declared no competing interest.

